# Hypoxia differentially affects coronary vessel formation during heart development

**DOI:** 10.64898/2026.02.06.704033

**Authors:** Sophie Payne, Susann Bruche, Dorota Szumska, Alice Neal, Mark D Preston, Sarah De Val

## Abstract

**BACKGROUND:** The coronary vessel system is a dense and diverse network of arteries, veins and capillaries formed by endothelial cells from a variety of sources. While hypoxia is a known stimulus for angiogenic sprouting generally, the exact mechanisms by which hypoxia, and resultant increased VEGFA, influences vessel growth in the heart are not clearly delineated.

**METHODS:** We used a genetic model to mimic hypoxia through ectopic stabilisation of myocardial HIFα. This enabled us to study the consequences of hypoxia without vascular depletion. Changes in coronary ECs in these hearts relative to littermate controls were assessed by single cell RNA-sequencing, and by examining the activity of enhancer:reporter transgenes active in different coronary vessel beds downstream of distinct vascular regulatory pathways.

**RESULTS:** Analysis of hypoxia-mimic hearts found increased angiogenic gene expression alongside expanded activity of the VEGFA-MEF2-driven angiogenic regulatory pathway in a pattern that indicated increased endocardial-derived angiogenic sprouting. Conversely, regulatory pathways specifically active in the sinus venosus (SV)-derived plexus showed little variance in response to stabilized HIFα, and sprouting from the SV was not expanded. Although hypoxia and increased VEGFA levels have been previously linked to increased arterial differentiation, we saw little change in initial arterial EC differentiation in the experimental hearts. However, mature coronary arterial formation was delayed.

**CONCLUSIONS:** These observations further emphasize a direct and specific link between hypoxia and endocardial coronary vessel sprouting and suggest a role of hypoxia/VEGFA in guiding coronary arterial coalescence.

## INTRODUCTION

Endothelial cells (ECs) form the inner layer of all blood vessels and are the first component of the vascular system to form. Whilst the initial endothelial plexus in the early embryo is established by EC differentiation from mesodermal progenitors (vasculogenesis), most subsequent growth of new blood vessels occurs from existing ones, a process known as angiogenesis. During sprouting angiogenesis, the most intensely investigated form of angiogenesis, ECs take on a filopodia-rich migratory tip cell phenotype and invade the interstitial space, followed by highly-proliferative ECs known as stalk cells ^1–3^. Sprouting angiogenesis can be induced by hypoxic conditions and is stimulated by growth factors such as VEGFA (via VEGFR2) and chemokines such as CXCL12 (via CXCR4) ^1–3^.

Although initially avascular, the high metabolic demands of the heart necessitate the formation of a coronary vasculature during embryonic development, a process that begins on embryonic day 11 (E11) in mice. Coronary vessels are formed by ECs from existing vascular structures and therefore grow via angiogenesis by definition. However, the ECs within the coronary vasculature come from multiple developmental origins, with the endocardium and sinus venosus (SV) representing the two main sources ^4–7^. The endocardium, an inner layer of specialised ECs within the heart, contributes to coronary vessels via endothelial sprouting outwards into the myocardium, a process which has been linked to hypoxia-induced myocardial VEGFA ^2,6^. Concurrently, venous ECs from the SV, a transient developmental structure at the venous pole of the heart, de-differentiate, sprout and migrate down the dorsal aspect of the heart to form a second coronary vascular plexus ^5,7,8^. However, although canonical Pdgfb-positive tip cells form during angiogenesis from both the endocardium and the subepicardial SV-origin plexus ^9^, SV-derived coronary sprouting is primarily linked to VEGFC and Elabela signalling rather than VEGFA ^5,8^. The developmental and functional heterogeneity of coronary ECs during development is also reflected by distinct expression profiles of some vascular gene (e.g. *Aplnr*/*Apj* in SV-derived vessels, *Fabp5* in endocardial-derived^10–12)^, and by differential activity of vascular enhancers (e.g. enhancers independently activated by VEGFA-MEF2, SOXF/RBPJ and BMP-SMAD1/5 are expressed in different compartments of the coronary vasculature^13^). However, despite their independent origins and stimuli, endocardial-derived and SV-derived coronary networks share expression of some angiogenic-associated genes and ECs from both origins contribute to coronary artery and vein formation ^9,12,14^. Additionally, these two vascular networks merge at late fetal stages to ensure the entire heart is vascularised, and ECs from the two origins are genetically and functionally indistinguishable from each other in the adult heart ^15^.

It has been hypothesised that endocardial-derived coronary vascularization is selectively physiologically responsive to microenvironmental cues including hypoxia, whilst SV-derived coronary vascularization is instead developmental timed and genetically driven ^8^. This grew out of observations that endocardial-derived vessels compensate for the loss of SV-derived vasculature during embryonic development, a redundancy attributed to increased myocardial hypoxia and subsequent VEGFA expression ^8^. However, these experiments were unable to directly assess whether hypoxia had a similar ability to activate SV-derived coronaries, as these were depleted in order to drive hypoxia in the first place. Additionally, coronary vessel growth in the adult heart, closely associated with hypoxia, does not originate from endocardial ECs but instead predominantly involves angiogenesis from established coronary vessels within the myocardial wall ^16^. It is therefore unclear whether this occurs via the same pathways involved in embryonic coronary formation. This is particularly important because the angiogenic response to myocardial infarction and ischemic injuries in the adult mammalian heart is limited and insufficient. Efforts to improve new vessel growth therapeutically have focused on reactivating canonical angiogenic pathways such as VEGFA-VEGFR2 but have shown little success ^17^, with the responsiveness of coronary ECs to pro-angiogenic signals appearing reduced compared to ECs in other tissues ^18^.

Here, we investigate the consequences of a hypoxic environment on the transcriptome of coronary ECs in the developing heart, and on different aspects of vascular growth in the heart as it becomes vascularized. This analysis takes advantage of the spatially distinct nature of coronary vascular development, allowing us to determine the effects of myocardial hypoxia on sprouting angiogenesis from different sources and driven by different pathways, and on arteriovenous differentiation and maturation.

## METHODS

### Animals

All animal procedures were approved by a local ethical review committee at Oxford University and licensed by the UK Home Office and follow ARRIVE guidelines. The published mouse alleles used were: *Phd2^flox/flox^*^19^; Nkx2.5-Cre ^20^; Rosa26-*tdTomato*^21^; ROSA26-*LacZ* ^22^; HLX-3:*LacZ* ^23^, NOTCH1+16:*LacZ* ^24^, Dll4-12:*LacZ* ^25^ and Ephb4-2:*LacZ* ^26^. All mice were back-crossed at least 5 generations onto a pure C57BL/6J background.

For embryo collection, matings were set up and female mice checked for vaginal plugs every morning. The date of observation of vaginal plug was considered embryonic day (E)0.5. Embryos were harvested into phosphate buffered saline (PBS) on ice, embryonic hearts then dissected and processed for X-gal staining, cryo-embedding in OCT Embedding Medium (Thermo Scientific), or paraffin embedding. Analysis of enhancer:reporter activity was limited to litters including both control and experimental embryos, as the coronary vasculature grows quickly over a short period so small differences in developmental timepoint between different litters can impact analysis.

Genotyping for *Phd2^flox^*, *Cre* and *LacZ* was performed by PCR. Genomic DNA was extracted from ear biopsy (pups) or tail tip / yolk sac (embryos) samples by incubation for 1 hour at 98°C in 100µl 25mM NaOH / 0.2mM EDTA, followed by addition of 100µl 40mM Tris-HCl (pH=5.5). 1.5-2µl of this digestion mix was used directly for the PCR reaction.

### Single-cell RNA sequencing

*Phd2^fl/fl^;*Rosa26-*tdTomato^hom^* adult female mice were mated with *Phd2^fl/fl^;Nkx2.5-Cre^het^* males and plug checked each morning. Four *Phd2^fl/fl^* and four *Phd2^fl/fl^;Nkx2.5-Cre* embryos were harvested at E13.5 from a single litter, and hearts dissected on ice with outflow tracts and atria removed. Fluorescent microscopy was used to distinguish between control *Phd2^fl/fl^*;Rosa26-*tdTomato^het^*and *Phd2^fl/fl^;Nkx2.5-Cre*;Rosa26-*tdTomato^het^*hearts, with all genotypes later confirmed by PCR on tail tips. Each heart was processed as an individual sample for the scRNA-seq experiment, and all samples were processed together to minimise batch effects in the bioinformatic analysis.

For enzymatic and mechanical dissociation of the hearts, the Neonatal Heart Dissociation Kit (Miltenyi Biotec, 130-098-373) was used according to manufacturer’s instructions in combination with the gentleMACS^TM^ Octo Dissociator with Heaters (Miltenyi Biotec), including the red blood cell lysis steps. The resulting cell pellets were then resuspended in 100µl of 3% FBS in PBS containing 1:100 anti-mouse-CD31 conjugated to FITC (BioLegend, 102406) and incubated on ice for 25 minutes. 100µl of 1µg/ml DAPI solution was then added to each sample and incubated on ice for a further 5 minutes before cell suspensions were centrifuged at 600x g for 5 minutes at 10°C. Cell pellets were then resuspended in 500µl of 2% FBS in PBS, transferred to FACS tubes and kept on ice in the dark. Single-stained and unstained cells were also prepared as controls for setting up the FACS machine.

Cells were sorted using a Sony MA900 FACS Sorter, with gating set up first using an FSC-A vs BSC-A plot to exclude cell debris, then an FSC-A vs FSC-H plot for the exclusion of doublet cells. Cell viability was determined using DAPI staining, with low DAPI cells designated as ‘Live’ cells, and unstained / single-stained samples used to determine a suitable threshold for CD31-FITC staining. CD31-positive ECs from each sample were sorted into 100µl 3% FBS in PBS. The FACS cell counts ranged from 5919 to 10580 per heart, and when all samples had been run on the Sorter the collected cells were centrifuged at 600x g for 5 minutes at 10°C, then the supernatant was aspirated, and the cell pellets resuspended in 30µl 0.04% BSA in PBS.

Samples were then immediately processed for sequencing using the Chromium Next GEM Single Cell 3’ GEM, Library & Gel Bead Kit v3.1 from 10X Genomics according to manufacturer’s instructions. Each sample was processed in separate GEM well, and then samples were pooled to a single flow cell and run on a NextSeq2000 sequencer. The raw read data was downloaded from Illumina BaseSpace for quality control and bioinformatic analysis.

### Bioinformatic analysis

We followed the standard Cell Ranger pipelines to demultiplex the raw base call (BCL) files, align reads to the mouse mm10 reference genome, generate feature-barcode matrices and aggregate the samples ^27^. The resulting sequencing and alignment statistics were used as quality controls (Supplemental Figure S3A-B). The dataset was used to create a SeuratObject in R for further processing and analysis: following normalisation across samples, cells were excluded if they had fewer than 200 or more than 7000 genes, contained more than 10% mitochondrial RNA, or were assessed as doublets using DoubletFinder^28^. Datasets were projected into 2-dimensional space using the uniform manifold approximation and projection (UMAP) algorithm, followed by unsupervised graph-based clustering.

Cluster annotation was performed manually based on the most upregulated genes per cluster and previously reported marker genes. Cell cycle status between control and “hypoxic” cells was assigned using the Seurat CellCycleScoring() function, and differentially expressed genes were assigned using the FindMarker() function with default Wilcoxon Rank Sum test. The gprofiler2 package was used for pathway analysis on genes upregulated (p_val_adj < 0.05 and avg_log2FC > 0.2) in all cells in hypoxic conditions.

For analysis on coronary ECs only, clusters 9 and 15 were selected as a subset before repeating UMAP projection, graph-based clustering and annotation. For lists of top 10 up/downregulated genes per cluster ranked by p value / log2FC, only genes with a value of >0.1 in both pct.1 and pct.2 were included.

### Wholemount Immunostaining

After dissection in cold PBS, embryonic hearts were fixed for 30 minutes (E12.5 hearts) or 40 minutes (E13.5 hearts) in 4% paraformaldehyde (PFA) in PBS at 4°C, before dehydration through a methanol/PBS series to 100% methanol and storage at −20°C. For immunostaining, hearts were rehydrated back through the methanol series to PBS, permeabilised for 1 hour in 1% Triton-X100 in PBS at room temperature, and then incubated with primary antibodies diluted in 0.5% Triton-X100 in PBS at 4° for 1-4 days. Primary antibodies used were Armenian hamster anti-CD31 (Abcam, ab119341, diluted 1:200), rat anti-Endomucin (Santa Cruz, sc-65495, diluted 1:50), and goat anti-SOX17 (R&D, AF1924, diluted 1:500). After primary antibody incubation, hearts were washed 5x 30 minutes in PBS, followed by overnight incubation at 4°C with Alexa Fluor®-conjugated secondary antibodies diluted 1:500 in 0.2% TritonX-100 in PBS. Hearts were finally washed for a further 5x 30 minutes in PBS, fixed in 4% PFA in PBS for 30 minutes, rinsed twice in PBS and then mounted onto concave cavity microscope slides with Fluoromount^TM^ aqueous mounting medium (Sigma-Aldrich). Confocal stacks were obtained using a Zeiss 710 MP confocal microscope and processed using Zen and ImageJ software.

### Whole-mount X-gal Staining

β-galactosidase expression from the *LacZ* gene was detected by X-gal staining. Following dissection of embryonic hearts in cold PBS, samples were fixed in 2% PFA / 0.2% glutaraldehyde in PBS at 4°C for 30-60 minutes depending on embryonic stage. They were then washed twice for 20-30 minutes in Rinse solution (2mM MgCl_2_, 0.2% Nonidet P40, 0.1% sodium deoxycholate in PBS) before incubation in Staining solution (Rinse solution containing 1mg/ml 5-bromo-4-chloro-3-indolyl β-d-galactopyranoside (X-gal), 5mM K_4_Fe(CN)_6_ and 5mM K_3_Fe(CN)_6_). For visualising the HLX-3:*LacZ* and Dll4-12:*LacZ* transgenes, hearts were stained at room temperature for 3 hours, and for visualising the NOTCH1+16:*LacZ*, EphB4-2:*LacZ* and Rosa26-*LacZ* Cre reporter samples were incubated in the Stain solution overnight. Imaging of whole hearts was performed using a stereo microscope (Leica M165C) equipped with a ProgRes CF Scan camera and ProgRes CapturePro software (Jenoptik). Only *Phd2^fl/fl^;Nkx2.5-Cre* hearts carrying one of the enhancer:*LacZ* transgenes with at least one *LacZ*-positive *Phd2^fl/fl^* control littermate were included in analysis, due to small changes in embryonic stage reached between litters and the rapid development the coronary vasculature.

Following X-gal staining, some hearts and embryos were dehydrated through an ethanol series, cleared using Histo-Clear (National Diagnostics) and paraffin-embedded for sectioning. 10μm sections were de-waxed, counter-stained using Nuclear Fast Red (Electron Microscopy Services) and imaged using a NanoZoomer S210 slide scanner with NDP.view2 viewing software (Hamamatsu).

### X-gal Staining on Cryosections

X-gal staining was also performed directly on cryosections. Dissected E15.5 hearts were fixed for 45 minutes in 4% PFA in PBS at 4°C, then incubated overnight in 30% sucrose in PBS at 4°C. Following washes in a 50/50 mix of 30% sucrose/OCT Embedding Medium (Thermo Scientific) and then OCT, hearts were mounted in OCT over dry ice and stored at −80°C. Transverse cryosections were cut at a thickness of 10-15µm and stored at −80°C until ready to use.

Cryosections were thawed at room temperature before removal of the OCT by washing in PBS in a Coplin jar. Sections were then incubated in Fix solution (4% PFA, 2mM MgCl_2_, 5mM EGTA in PBS) for 10 minutes at room temperature, before further washes in PBS. They were then incubated in Cryosection Staining solution (2mM MgCl_2_, 0.02% Nonidet P40, 0.01% sodium deoxycholate, 5mM K_4_Fe(CN)_6_, 5mM K_3_Fe(CN)_6_, and 1mg/ml X-gal in PBS) in a humidified chamber overnight at room temperature. Once the desired staining intensity was reached, sections were washed in PBS, fixed again in 4% PFA for 15 minutes, washed in PBS, and counter-stained with Nuclear Fast Red. After dehydration and mounting, slides were imaged using a NanoZoomer S210 slide scanner with NDP.view2 viewing software (Hamamatsu).

### *In situ* hybridisation

*In situ* hybridisation for detection of mRNAs was performed on paraffin sections using RNAScope® (PMID: 22166544). *Phd2^fl/fl^* and *Phd2^fl/fl^;Nkx2.5-Cre* littermate embryos at E13.5 and E15.5 were dissected in cold PBS and the chest region fixed overnight in 4% PFA in PBS at 4°C. They were then dehydrated through an ethanol series, cleared in Histo-Clear (National Diagnostics) and paraffin-embedded. Sections were cut at 6µm thickness, air dried overnight and stored at room temperature. *Phd2^fl/fl^*and *Phd2^fl/fl^;Nkx2.5-Cre* littermate hearts were processed together for each experiment comparing gene expression between the two genotypes.

The RNAscope® Multiplex Fluorescent Reagent Kit v2 (Bio-Techne Ltd) was used according to manufacturer’s instructions, with mild-standard conditions used for target retrieval (15 minutes in the Target Retrieval Reagent in a steamer) and protease digestion in the Protease Plus reagent for 30 minutes at room temperature. The probes used were: Mm-Vegfa-C3 (436961-C3) and Mm-Vegfc (492701), with Opal 570 and Opal 690 Dyes (Akoya Biosciences) used to detect C1 and C3 channels.

Immediately following the RNAScope protocol, slides were processed for immunohistochemical detection of Endomucin. Briefly, slides were washed twice in PBS before incubation in a humidified chamber at room temperature for 1 hour in Blocking solution (4% FBS, 10% donkey serum, 0.2% Triton X-100 in PBS). They were then incubated overnight at 4°C in rat anti-Endomucin antibody (Santa Cruz, sc-65495) diluted 1:50 in Blocking solution. After further washes and incubation for 1 hour in Alexa Fluor®488-conjugated anti-rat secondary antibody (Life Technologies) diluted 1:500 in Blocking solution, slides were counterstained with DAPI and mounted using Fluoromount^TM^ aqueous mounting medium (Sigma-Aldrich). Confocal images were obtained using a Zeiss 710 MP confocal microscope and processed using Zen and ImageJ software.

### Quantification of Imaging Data

For all quantification analysis, significance was tested using an unpaired t test, with p<0.05 determined as significant, and all N numbers are included in figure legends.

Ventricular wall thickness of Nuclear Fast Red stained transverse sections through E15.5 *Phd2^fl/fl^* and *Phd2^fl/fl^;Nkx2.5-CRE* hearts was measured using tools in the NDP.view2 software (Hamamatsu), with measurement at the most lateral point of the right and left ventricles.

Progression of the SV-derived vascular plexus on either CD31-immunostained or XGal-stained wholemount hearts at E12.5 and E13.5 was calculated manually in ImageJ, by measurement of the area from the SV to the furthest tips of the plexus as a percentage of the total area of the dorsal aspect of the ventricles.

Quantification of HLX-3 or NOTCH1+16 enhancer activity on the ventral aspect of E12.5 or E13.5 wholemount hearts was performed using a Macro in ImageJ to create a mask for blue XGal staining through colour analysis (based on methodology in ^29^) and measure its area, which was then converted to a percentage of the total area of the ventricles.

The same colour analysis pipeline was used to measure the area of blue XGal stain on E13.5 transverse sections through *Phd2^fl/fl^* and *Phd2^fl/fl^;Nkx2.5-CRE* hearts expressing the HLX-3:*LacZ* transgene, which again was calculated as a percentage of the total section area.

Quantification of Fluorescent RNAscope® results was performed in QuPath based on technical notes from ACD/Bio-Techne (QuPath Analysis Guidelines, MK 51-154/Rev A/Date 12/21/2020). Briefly, the ventricles were selected as the region of interest, and cell segmentation based on DAPI nuclear staining performed using an intensity threshold of 10 and the following nucleus parameters: Background radius 10µm, Median filter radius 2µm, Sigma 2µm, Minimum area 10µm^2^, Maximum are 300µm^2^. Vegfa or Vegfc probe detection was then performed using the subcellular detection module, with the following parameters that gave the best detection results: Expected spot size 0.5µm^2^, Minimum spot size 0.3µm^2^, Maximum spot size 2µm^2^. The percentage of cells containing at least one spot was calculated for each section and results compared across control and hypoxic genotypes.

## RESULTS

### Cardiomyocyte-specific knockdown of *Phd2* mimics hypoxia in the developing heart

Hypoxia inducible factors (HIF) are transcription factors that act as the master regulators of the cellular response to hypoxia. Under normal oxygen conditions, prolyl hydroxylase domain 2 containing protein (PHD2, encoded by the Phd2/*Egln1* gene) targets HIF-1α and HIF-2α for degradation ^30^. Here, we crossed mice homozygous for the conditional *Phd2^flox^* allele ^19^ with the constitutive *Nkx2.5-Cre* allele, which drives Cre expression in the cardiac crescent and cardiomyocytes from early in heart development prior to coronary vascular network formation (^20^ and Supplemental Figure S1). The subsequent loss of myocardial PHD2 ectopically stabilises HIFα proteins in the developing myocardium, inducing HIFα-dependent transcriptional programmes and genetically mimicking hypoxia (Figure 1A). *Phd2^fl/fl^;Nkx2.5-Cre* mice were viable and showed no overt phenotype. *Phd2^fl/fl^* females were crossed with *Phd2^fl/fl^;Nkx2.5-Cre* males, producing litters of control *Phd2^fl/fl^* and HIFα-stabilized, hypoxia-mimic *Phd2^fl/fl^;Nkx2.5-Cre* embryos (Figure 1B).

**Figure 1:**
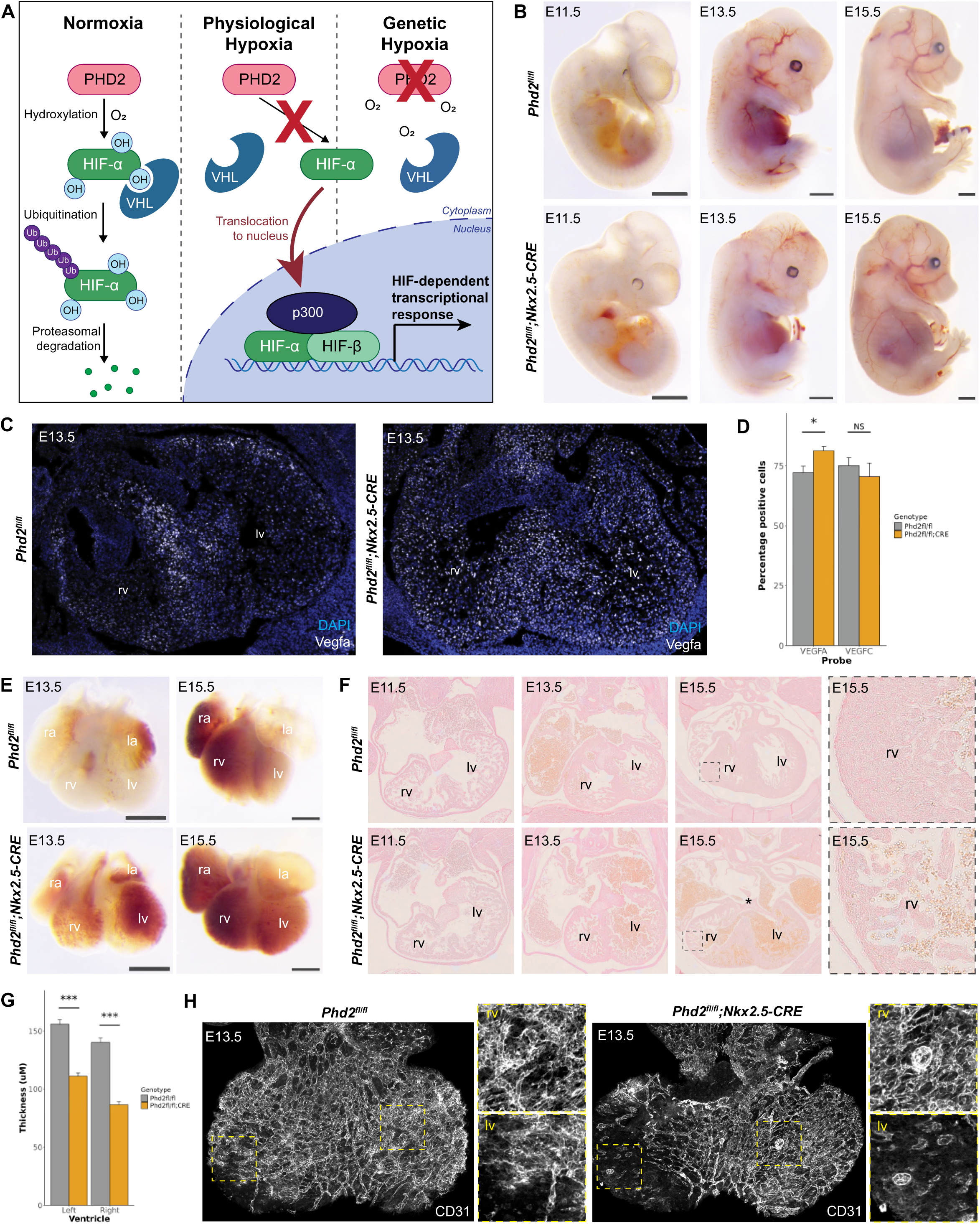
Myocardial-specific knockdown of *Phd2* **A** Schematic summarizing use of genetic knockdown of *Phd2* to mimic hypoxia. **B** Representative littermate *Phd2^fl/fl^* and *Phd2^fl/fl^;Nkx2.5-Cre* embryos at embryonic day (E)11.5, E13.5 and E15.5, scale bars = 1mm. **C** *In situ* hybridisation for *Vegfa* using RNAScope^TM^ on transverse sections through littermate E13.5 *Phd2^fl/fl^*and *Phd2^fl/fl^;Nkx2.5-Cre* hearts, representative of 3 repeats. **D** Quantification of *Vegfa* and *Vegfc* transcripts in E13.5 hearts recorded as percentage of cells in each section positive for each transcript. * = p<0.05, NS = not significant (unpaired *t* test), error bars show SEM, N=7 sections from 3 biological replicates per genotype. **E-G** Representative littermate *Phd2^fl/fl^* and *Phd2^fl/fl^;Nkx2.5-Cre* wholemount (**E**) or transversely sectioned hearts (**F-G**), scale bars = 0.5mm, asterisk marks a ventricular septal defect, dashed grey boxes indicate magnified images. Quantification of myocardial wall thickness (**G**) from N=28 sections through *Phd2^fl/fl^* heart and 50 sections through two *Phd2^fl/fl^;Nkx2.5-Cre* hearts in left and right ventricles at E15.5, *** = p<0.0001 (unpaired *t* test), error bars show SEM. **H** Dorsal views of representative E13.5 *Phd2^fl/fl^* and *Phd2^fl/fl^;Nkx2.5-Cre* hearts immunostained for CD31, yellow boxes indicate magnified views. Additional images in Supplemental Figure S2. rv = right ventricle; ra = right atrium; lv = left ventricle; la = left atrium.

Consistent with previous reports of hypoxia-induced increases in *Vegfa* transcription ^31,32^, loss of myocardial PHD2 resulted in a significant increase in *Vegfa* expression in *Phd2^fl/fl^;Nkx2.5-Cre* hearts comparative to controls, (Figure 1C-D). In control (Cre-negative) hearts, *Vegfa* expression was highest in the interventricular septum (IVS) and lower in the ventricles (Figure 1C), consistent with previous reports ^6^. Conversely, *Vegfa* expression in *Phd2^fl/fl^;Nkx2.5-Cre* hearts was more uniform across the whole myocardium (Figure 1C). *Vegfc* expression was not altered between *Phd2^fl/fl^* and *Phd2^fl/fl^;Nkx2.5-Cre* heart sections (Figure 1D)*. Phd2^fl/fl^;Nkx2.5-Cre* embryonic hearts also exhibited some morphological defects. *Phd2^fl/fl^;Nkx2.5-Cre* hearts collected at E11.5 and E13.5 showed no overall gross phenotype, but by E15.5 the overall shape of the hearts was altered, with a more rounded apex and enlarged atria (Figure 1E-F). Additionally, the ventricular walls were significantly thinner than controls, indicating a failure of compaction of the myocardium (Figure 1F-G), and membranous ventricular septal defect (VSD) were observed in two out of seven hearts (Figures 1F). CD31 staining of the vasculature showed blood islands on the dorsal side of the heart exclusively in *Phd2^fl/fl^;Nkx2.5-Cre* hearts at E12.5 (1/2) and E13.5 (2/2), and slightly reduced expansion of the SV-derived vascular plexus on the left ventricle at E13.5 (Figure 1H and Supplemental Figure S2 A, B and D). As expected, blood islands were also seen on the ventral aspect of all hearts at E13.5. Although there was a trend towards increased visible CD31-positive blood islands in *Phd2^fl/fl^;Nkx2.5-Cre* hearts (Supplemental Figure S2 C and E), this was non-significant.

### Single cell RNA-sequencing analysis of ECs in *Phd2^fl/fl^;Nkx2.5-Cre* hearts

Next, we performed single cell RNA-sequencing on ECs isolated from *Phd2^fl/fl^* and hypoxia-mimic *Phd2^fl/fl^;Nkx2.5-Cre* hearts at E13.5. Briefly, littermate hearts were dissected and Cre positive hearts identified through recombination of the *Rosa26-tdTomato* reporter allele (n=4 each of control and Cre-positive hearts). Each heart was processed as an individual sample for sequencing. Hearts were dissociated and CD31-positive ECs selected by FACS. Samples were then prepared for single cell RNA-sequencing using 10X Genomics Chromium Next GEM technology, and standard 10X Genomics bioinformatics analysis pipelines used to process the resulting reads, see Supplemental Figure S3A-B for sequencing statistics and QC metrics.

Cluster identification based on shared nearest neighbours resulted in 20 clusters (Figure 2A), 15 of which expressed common EC identity genes *(CD31, Kdr*) and were therefore assumed endothelial (Figure 2B). These EC clusters were split between endocardial (*Npr3* and *Adgrg6* positive) and coronary ECs (*Fabp4* and *Cldn5* positive) (Figure 2B): The four largest clusters (C0-3) all contained endocardial ECs, with high expression of markers for ventricular endocardium and transition to OFT, as defined by Cano et al. ^33^ (Figure 2C). A fifth endocardial EC cluster (C13) was located closer to the three clusters of valve ECs (C4, C10 and C16), and contained markers for EndoMT endocardium. A further three clusters of endocardial ECs were identified as ‘proliferating endocardium’ due to the high expression of G2M (C7 and C11) or S phase (C5) cell cycle markers. In addition, we identified two *Fabp4+*; *Cldn5*+ clusters of coronary EC clusters (C14-15), the smaller of which (C15) also expressed G2M markers (Figure 2C).

**Figure 2:**
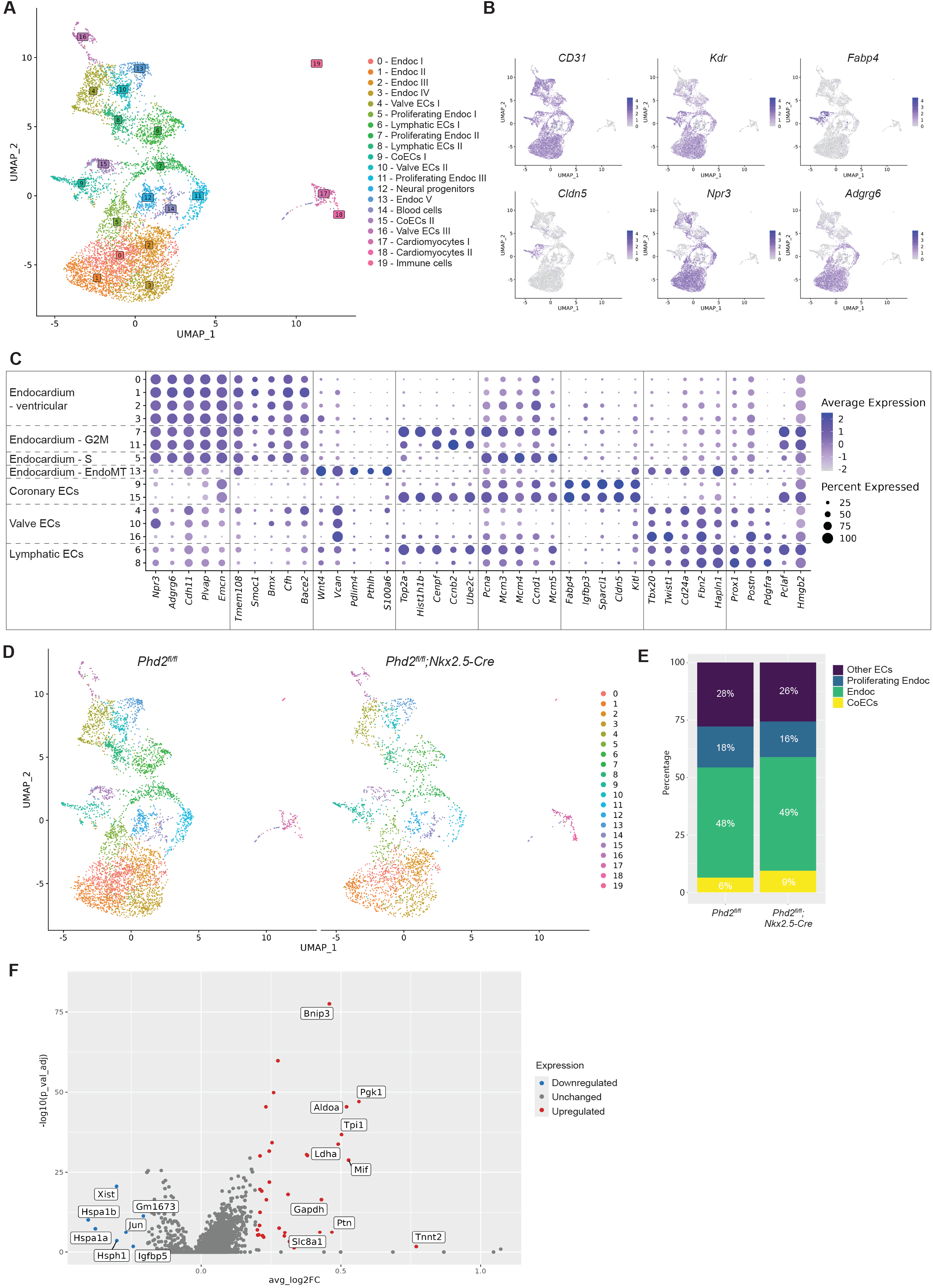
Single-cell sequencing analysis on ECs from *Phd2^fl/fl^* vs *Phd2^fl/fl^;Nkx2.5-Cre* hearts at E13.5 **A** UMAP projection of the scRNAseq dataset showing 20 clusters. **B** Expression of key markers of endothelium (*CD31*, *Kdr*), coronary endothelium (*Fabp4*, *Cldn5*), and endocardium (*Npr3*, *Adgrg6*) visualised on UMAP projections. **C** Dot blot to show key genes defining each sub-group of EC clusters, plotted with relative expression in each cluster. **D** Side-by-side comparison of UMAP projections for the cells from control *Phd2^fl/fl^* and “hypoxic” *Phd2^fl/fl^;Nkx2.5-Cre* hearts. **E** Bar chart to compare the percentage of cells in EC subtypes by genotype (coronary ECs (coECs) = clusters C9 and C15; Endocardium = C0, C1, C2, C3 and C13; Proliferating endocardium = C5, C11 and C13; Other ECs = C4, C6, C8, C10 and C16). n = 4336 *Phd2^fl/fl^* cells and 2400 *Phd2^fl/fl^;Nkx2.5-Cre* cells, percentage values rounded to nearest whole number. **F** Volcano plot to show up- and downregulated genes in *Phd2^fl/fl^;Nkx2.5-Cre* cells compared with *Phd2^fl/fl^* cells.

Side by side comparison of the UMAP projections for *Phd2^fl/fl^*and *Phd2^fl/fl^;Nkx2.5-Cre* genotypes showed very similar distribution of cells across the clusters (Figure 2D), indicating that hypoxic myocardium neither results in the loss of an EC subtype or cell state, nor creates a new one. Dividing the EC clusters into broader groups representing endocardium, proliferating endocardium, coronary ECs and other ECs, we saw an increase of coronary ECs within the EC group in *Phd2^fl/fl^;Nkx2.5-Cre* hearts, rising approximately 50% from 6% to 9% (Figure 2E). Differential gene expression analysis across all ECs identified 45 differentially expressed genes (DEGs) in *Phd2^fl/fl^;Nkx2.5-Cre* cells compared with *Phd2^fl/fl^*cells (adjusted p-value after multiple testing correction < 0.05 and log2 fold change > 0.2, Figure 2F and Supplemental Table 1). Of these, 37 genes were significantly upregulated in the hypoxia-like hearts, whilst 8 genes were significantly downregulated. Pathway analysis on the upregulated genes produced many terms associated with metabolic processes, particularly relating to glycolysis and carbon metabolism (Supplemental Table 2).

### Coronary ECs shift towards pro-angiogenic tip cell and non-cycling populations in *Phd2^fl/fl^;Nkx2.5-Cre* hearts

To focus in on the coronary ECs, we selected the cells from clusters C9 and C15, and re-ran the UMAP generation and unsupervised clustering (see Supplemental Figure S3C for QC metrics). There were 276 *Phd2^fl/fl^* and 225 *Phd2^fl/fl^;Nkx2.5-Cre* cells, which divided into 7 clusters (Figure 3A), and were positive for endothelial and coronary markers (Figure 3C). The four largest clusters were annotated as capillary coronary ECs, with cluster C0 showing G1 markers and clusters C1-3 showing cell cycle markers suggesting they are actively proliferating (Figure 3D). This reflects the relatively undifferentiated state of the coronary vasculature at E13.5, although smaller clusters were also identified containing *Gja5* and *Cxcr4*-positive pre-arterial cells, *Igfbp3* and *Kcne3*-positive tip cells, and *Nr2f2*-positive venous ECs, consistent with previously reported cell types at this stage (Figure 3C and D).

**Figure 3:**
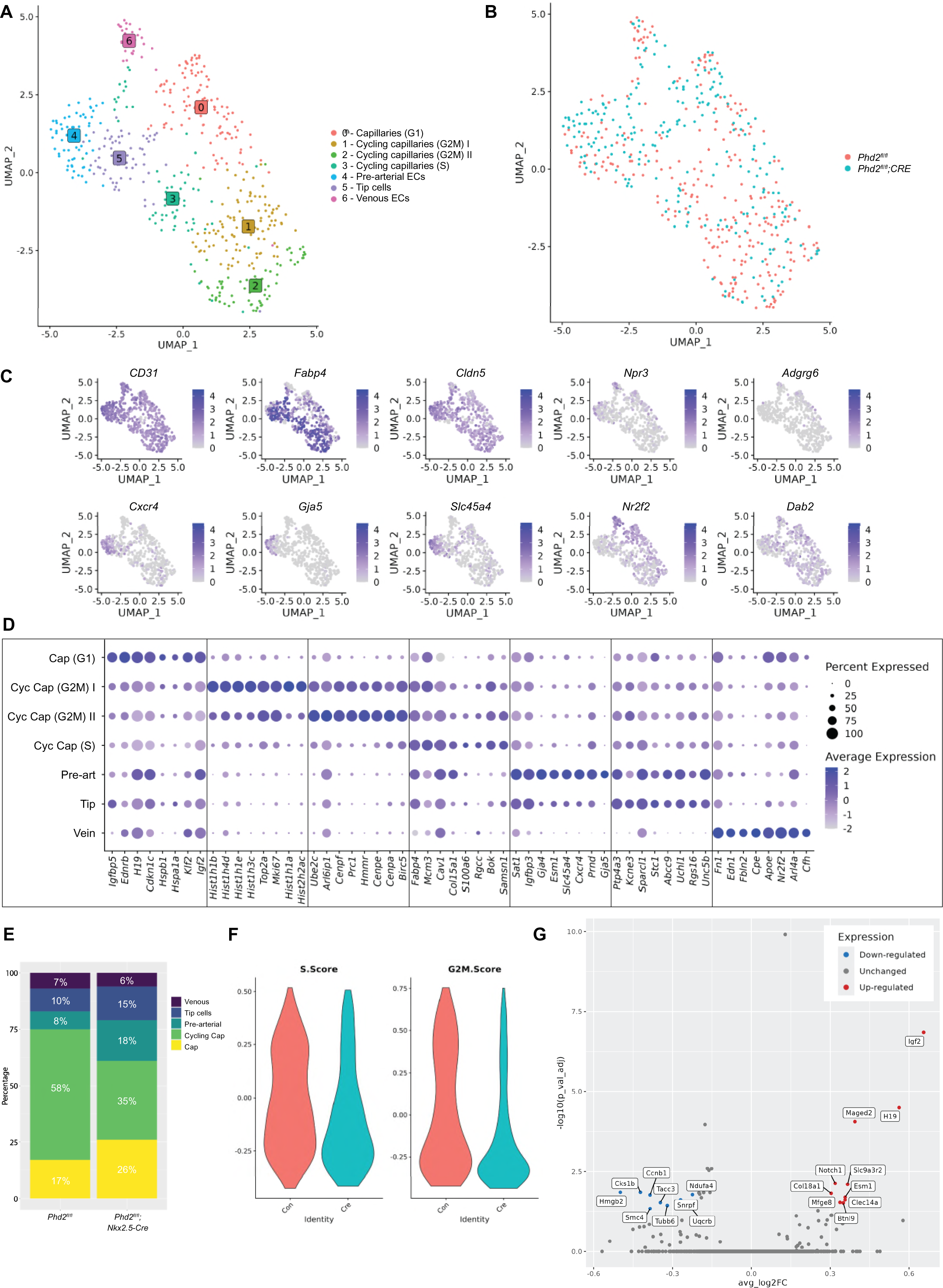
Single-cell sequencing analysis on coronary ECs only **A** Re-running of UMAP projection and clustering of coronary EC cells resulted in 7 clusters. **B** Overlay comparison of UMAP projections for cells from *Phd2^fl/fl^* and *Phd2^fl/fl^;Nkx2.5-Cre* hearts. **C** Expression of key markers of endothelium (*CD31*), coronary endothelium (*Fabp4*, *Cldn5*), endocardium (*Npr3*, *Adgrg6*), arterial (*Cxcr4*, *Gja5*, *Slc45a4*) and venous (*Nr2f2*, *Dab2*) genes, visualised on UMAP projections. **D** Dot blot to show key genes defining each sub-group of coEC clusters, plotted with relative expression in each cluster. **E** Bar chart to compare the percentage of cells classified as G1 capillaries (Cap), G2/M/S cycling capillaries (Cycling cap), pre-arterial cells, tip cells and venous cells between *Phd2^fl/fl^* and *Phd2^fl/fl^;Nkx2.5-Cre* hearts. Percentage values rounded to nearest whole number. **F** Violin plots showing S and G2M cell cycle scores assigned to each cell from *Phd2^fl/fl^* (red) and *Phd2^fl/fl^;Nkx2.5-Cre* (blue) hearts. **G** Volcano plot to show up- and downregulated genes in *Phd2^fl/fl^;Nkx2.5-Cre* cells compared with *Phd2^fl/fl^* cells.

Overlaying the UMAP distributions of the coronary ECs from the *Phd2^fl/fl^* and *Phd2^fl/fl^;Nkx2.5-Cre* hearts again indicated that all clusters contained cells from both genotypes (Figure 3B). However, there was a drop in the proportion of cycling capillaries in *Phd2^fl/fl^;Nkx2.5-Cre* hearts (35%, compared with 58% of control), with an accompanying increase in non-cycling capillaries (26% compared with 17%) (Figure 3E). A greater proportion of coronary ECs were also assigned as tip cells and pre-arterial cells in the hypoxic hearts, whilst the proportion of venous cells remained similar. The differences in coronary ECs between *Phd2^fl/fl^*and *Phd2^fl/fl^;Nkx2.5-Cre* hearts was also reflected in cell cycle analysis of the cohorts: there is an overall decrease in the S and G2M scores of coronary ECs from the *Phd2^fl/fl^;Nkx2.5-Cre* hearts, consistent with the drop in cycling capillaries (Figure 3F).

Differential gene expression analysis of all coronary ECs identified 18 DEGs in the coronary ECs exposed to hypoxic myocardium that passed multiple testing corrections (Table 1, Figure 3G). The 10 up-regulated genes include genes linked with angiogenesis, EC migration and tube formation, such as *Esm1*, *Igf2*, *H19* and *Clec14a* ^34^. Due to relatively small cell numbers, the same differential gene expression analysis on the individual EC clusters did not result in any significant DEGs after multiple testing correction. However, analysis of most differentially altered genes (ranked by log2 fold change and p value) from each cluster suggested an increase in angiogenic-associated genes, particularly in the cycling capillaries and venous EC clusters, suggesting it is these coronary EC subtypes that are most likely to undergo transcriptional changes towards an angiogenic phenotype in response to environmental hypoxia. (Supplemental Table 3).

**Table 1:** Differential gene expression analysis on all coronary ECs at E13.5 Differential gene expression analysis results comparing all *Phd2^fl/fl^;Nkx2.5-Cre* with control *Phd2^fl/fl^*cells, showing 10 genes upregulated in *Phd2^fl/fl^;Nkx2.5-Cre* cells and 8 genes downregulated in *Phd2^fl/fl^;Nkx2.5-Cre* coronary ECs. Genes were included if adjusted p-value after multiple testing correction < 0.05 and log2 fold change > 0.2.

| Gene | p_val | avg_log2FC | p_val_adj | Direction |
| --- | --- | --- | --- | --- |
| ***Igf2*** | 4.38E-12 | 0.655456846 | 1.41E-07 | Upregulated |
| ***H19*** | 9.71E-10 | 0.562078504 | 3.14E-05 | Upregulated |
| ***Maged2*** | 2.71E-09 | 0.393790579 | 8.75E-05 | Upregulated |
| ***Slc9a3r2*** | 2.48E-07 | 0.366018414 | 0.007991 | Upregulated |
| ***Esm1*** | 6.21E-07 | 0.356745856 | 0.020056 | Upregulated |
| ***Clec14a*** | 7.53E-07 | 0.356505389 | 0.024316 | Upregulated |
| ***Btnl9*** | 9.40E-07 | 0.349411029 | 0.030341 | Upregulated |
| ***Mfge8*** | 9.06E-07 | 0.337202444 | 0.029235 | Upregulated |
| ***Notch1*** | 2.32E-07 | 0.318960456 | 0.00749 | Upregulated |
| ***Col18a1*** | 4.73E-07 | 0.30355086 | 0.015269 | Upregulated |
| ***Snrpf*** | 7.49E-07 | -0.269404535 | 0.024175 | Downregulated |
| ***Tubb6*** | 1.13E-06 | -0.319949953 | 0.036531 | Downregulated |
| ***Tacc3*** | 9.20E-07 | -0.346286681 | 0.029715 | Downregulated |
| ***Smc4*** | 1.42E-06 | -0.38533039 | 0.045805 | Downregulated |
| ***Ccnb1*** | 5.31E-07 | -0.385563554 | 0.017149 | Downregulated |
| ***Cks1b*** | 4.38E-07 | -0.42225002 | 0.014144 | Downregulated |
| ***Hmgb2*** | 4.38E-07 | -0.49867428 | 0.014147 | Downregulated |
| ***Rps2*** | 2.28E-40 | -0.615811604 | 7.36E-36 | Downregulated |

In conclusion, this scRNAseq analysis suggested an increase in angiogenic gene expression in coronary ECs in response to a hypoxic environment, with a shift towards the non-proliferative, migratory tip cell EC phenotype.

### Hypoxic myocardium results in expanded and ectopic activation of the VEGFA-MEF2-dependent angiogenic pathway but does not affect SV-derived plexus formation

Our results so far indicate an increase in angiogenic ECs within the coronary vasculature in response to hypoxia-like HIFα stabilization. These observations align with similar accelerated coronary vessel growth reported in hearts after myocardial deletion of VHL, an alternative method of mimicking hypoxia^35^. We next investigated whether the increased angiogenic ECs in *Phd2^fl/fl^;Nkx2.5-Cre* embryonic hearts were specific to endocardial-derived sprouting or instead represented an increase in angiogenesis from all coronary origins. Additionally, we sought to establish the vascular regulatory pathways targeted by these hypoxic-like conditions.

Our previous work on coronary vascular regulatory pathways identified two transcriptional pathways active during the sprouting of the coronary vasculature, differentially driven by MEF2 and SOXF factors ^13^. The MEF2 transcriptional pathway is downstream of VEGFA and is linked to expression of angiogenic genes such as *Hlx* and *Dll4* in the systemic vasculature, where it is specifically active in angiogenic ECs ^13^. In the developing embryonic heart, enhancers regulated by this VEGFA-MEF2 pathway are predominately active in ECs sprouting from the endocardium (Figure 4A and ^13^). Conversely, enhancers regulated by the SOXF pathway are predominantly active throughout the SV-derived endothelial plexus (Figure 4A and ^13^). Whilst these patterns of expression align with the concept of distinct VEGFA-driven endocardial sprouting (MEF2) and VEGFC/Elabella-driven SV-derived sprouting (SOXF), the activity patterns of enhancers downstream of these two pathways already hints at some level of greater complexity: VEGFA-MEF2-driven enhancers are also transiently active at the leading edge of the SV-derived plexus as it initially moves over the dorsal ventricle between E11.5-E13.5 (Figure 4 and ^13^), suggesting that the classical angiogenic VEGFA-MEF2 pathways may be involved to some degree in both endocardial-derived and SV-derived sprouting. Conversely, whilst SOXF-driven enhancers are active throughout the SV-derived plexus as it sprouts down the dorsal side of the heart between E11.5-E13.5, they are also expressed in a subset of ECs emerging from the endocardium/septum region on the ventral side (Figure 4A, Figure 5 and ^13^). This expression pattern closely mimics that of SOX17, which is widespread throughout the SV-derived vascular plexus but also in active ECs sprouting from the endocardium at these timepoints (Supplemental Figure S4 and ^8,12^), again suggesting that angiogenic sprouting from both endocardial and SV origins may incorporate multiple inputs via different regulatory pathways.

**Figure 4:**
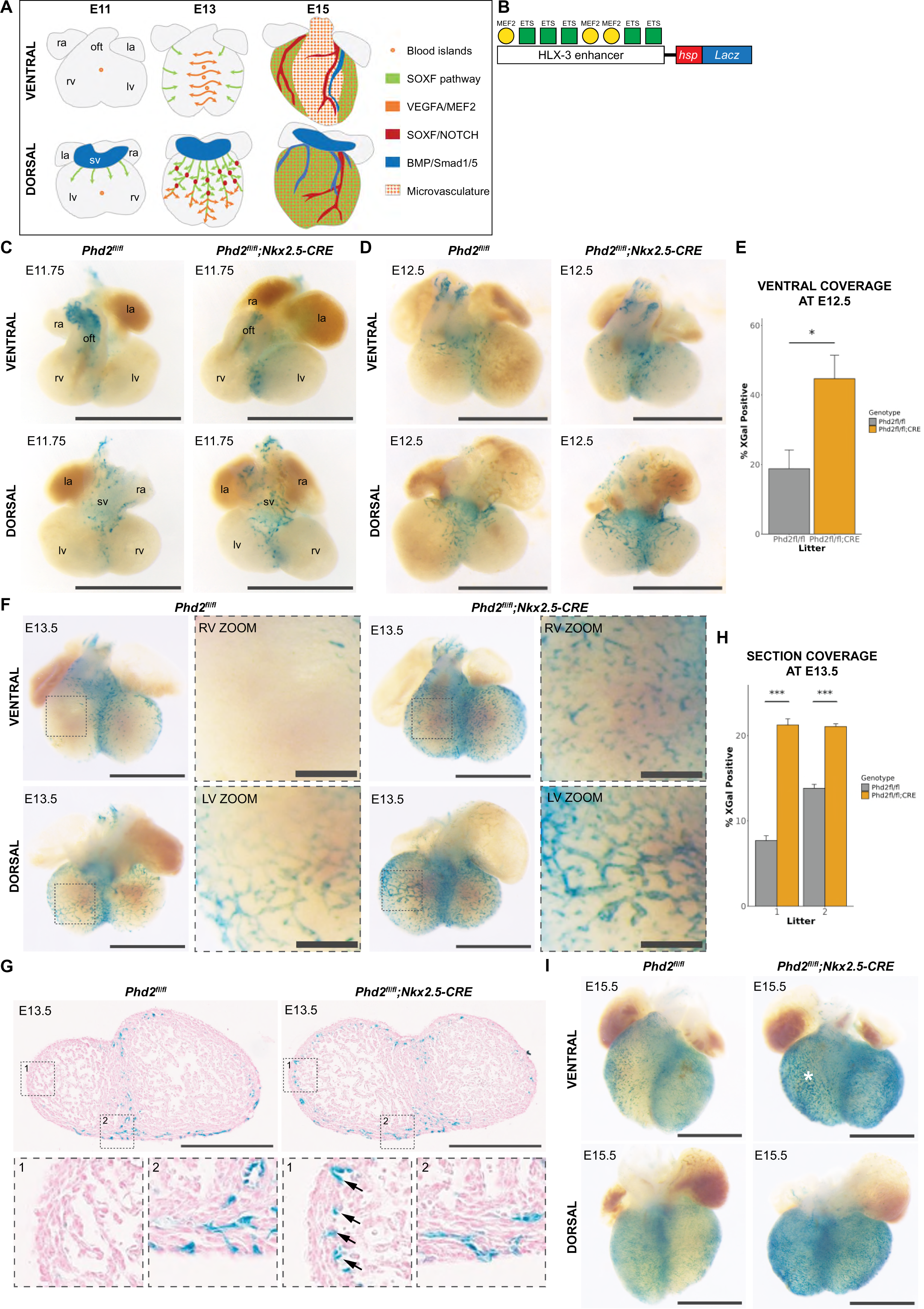
Expansion and ectopic activity of MEF2-dependent angiogenic pathway in hypoxic hearts **A** Summary of regulatory pathways driving coronary vascular growth E11.5-E15.5 ^13^. **B** Schematic of the HLX-3:*LacZ* transgene, with the human HLX-3 enhancer sequence cloned upstream of the hsp68 minimal promoter and *LacZ* reporter gene. Validated ETS and MEF2 TF binding sites within the enhancer are represented by coloured circles and squares ^23^. **C** Representative ventral and dorsal views of E11.75 littermate hearts with HLX-3:*LacZ* activity (blue XGal stain) on *Phd2^fl/fl^* and *Phd2^fl/fl^;Nkx2.5-CRE* genetic backgrounds. N= 3 *Phd2^fl/fl^;Nkx2.5-CRE* hearts, all other hearts shown in Supplemental Figure S5A. Scale bars = 1mm. ra = right atrium; la = left atrium; rv = right ventricle; lv = left ventricle; sv = sinus venosus; oft = outflow tract. **D** Representative E12.5 littermate hearts with HLX-3:*LacZ* activity on *Phd2^fl/fl^* and *Phd2^fl/fl^;Nkx2.5-CRE* genetic backgrounds. N=5 *Phd2^fl/fl^;Nkx2.5-CRE* hearts, all other hearts in Supplemental Figure S5B. Scale bars = 1mm. **E** Quantification of ventral area covered by HLX-3:*LacZ* activity on all hearts collected at E12.5, calculated as a percentage of the total ventral area. * = p<0.05 (unpaired *t* test), error bars show SEM, N=5 *Phd2^fl/fl^;Nkx2.5-CRE* hearts and N=4 *Phd2^fl/fl^* hearts. **F** Representative E13.5 littermate *Phd2^fl/fl^* and *Phd2^fl/fl^;Nkx2.5-CRE* hearts with HLX-3:*LacZ* activity, with magnified views of the RV on ventral side, and LV on dorsal side, highlighting ectopic HLX-3:*LacZ* activity in *Phd2^fl/fl^;Nkx2.5-CRE* hearts. N=7 *Phd2^fl/fl^;Nkx2.5-CRE* hearts, all other hearts shown in Supplemental Figure S6. Scale bars = 1mm in wholemount views and 0.2mm in magnified images. **G** Transverse sections through E13.5 *Phd2^fl/fl^* and *Phd2^fl/fl^;Nkx2.5-CRE* hearts, showing blue HLX-3:*LacZ* activity. Zoom 1 illustrates ectopic endocardial activity in *Phd2^fl/fl^;Nkx2.5-CRE* hearts highlighted with black arrows, Zoom 2 shows similar XGal-positive vessels migrating from the outer heart surface into the myocardium. Scale bars = 0.5mm. **H** Quantification of HLX-3:*LacZ* activity in two pairs of littermate E13.5 hearts, measuring the XGal-positive area as a percentage of total area of transverse sections. *** = p<0.0001 (unpaired *t* test), error bars show SEM, N=42 and 48 sections through *Phd2^fl/fl^* and *Phd2^fl/fl^;Nkx2.5-CRE* hearts respectively in litter 1, and N=41 and 31 sections through *Phd2^fl/fl^* and *Phd2^fl/fl^;Nkx2.5-CRE* hearts in litter 2. **I** Representative E15.5 littermate *Phd2^fl/fl^* and *Phd2^fl/fl^;Nkx2.5-CRE* hearts with HLX-3:*LacZ* activity, white asterisk highlights ectopic transgene activity on the ventral RV. N=7 *Phd2^fl/fl^;Nkx2.5-CRE* hearts, all other hearts shown in Supplemental Figure S7, including 3 *Phd2^fl/+^;Nkx2.5-CRE* hearts as CRE controls. Scale bars = 1mm in wholemount views and 0.2mm in magnified images. rv = right ventricle; ra = right atrium; lv = left ventricle; la = left atrium; sv = sinus venosus; oft = outflow tract.

**Figure 5:**
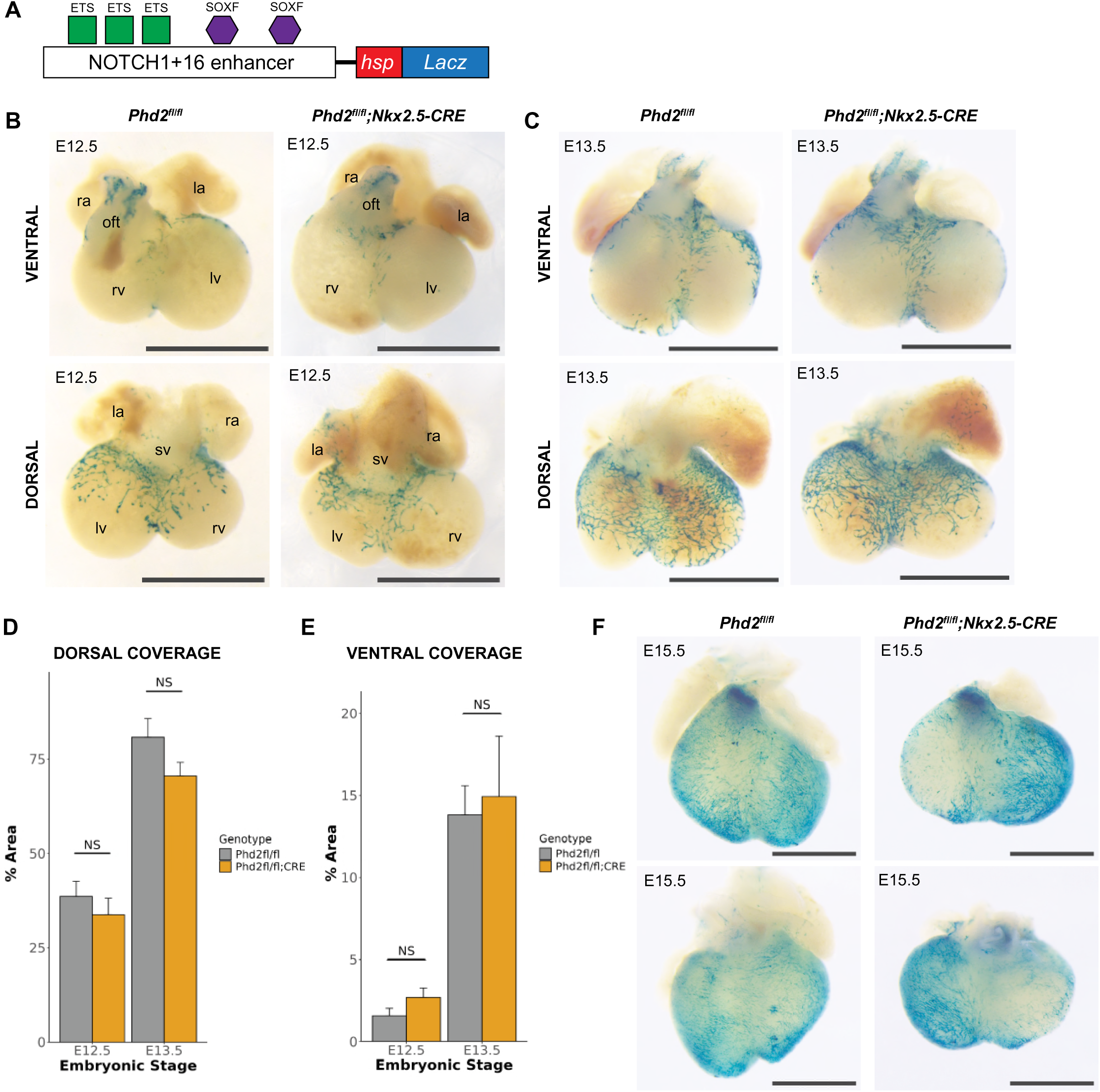
SV-derived vascular plexus formation is largely unaffected by hypoxic myocardium **A** Schematic of the NOTCH1+16:*LacZ* transgene, with human NOTCH1+16 enhancer sequence cloned upstream of the hsp68 minimal promoter and *LacZ* reporter gene. Validated ETS and SOXF TF binding sites within the enhancer are represented by coloured shapes ^24^. **B** Representative ventral and dorsal views of E12.5 littermate hearts with NOTCH1+16:*LacZ* activity (blue XGal stain) on *Phd2^fl/fl^* and *Phd2^fl/fl^;Nkx2.5-CRE* genetic backgrounds. N= 7 *Phd2^fl/fl^;Nkx2.5-CRE* hearts, all other hearts shown in Supplemental Figure S8A. Scale bars = 1mm. ra = right atrium; la = left atrium; rv = right ventricle; lv = left ventricle; sv = sinus venosus; oft = outflow tract. **C** Representative E13.5 *Phd2^fl/fl^* and *Phd2^fl/fl^;Nkx2.5-CRE* hearts showing NOTCH1+16:*LacZ* activity. N= 5 *Phd2^fl/fl^;Nkx2.5-CRE* hearts, all other hearts shown in Supplemental Figure S8B. Scale bars = 1mm. **D** Quantification of the extension of the NOTCH1+16:*LacZ*-positive SV-derived vascular plexus as a percentage of the total dorsal area of *Phd2^fl/fl^*and *Phd2^fl/fl^;Nkx2.5-CRE* hearts at E12.5 and E13.5 hearts. NS = not significant (unpaired *t* test), error bars show SEM, N=9 control and 7 *Phd2^fl/fl^;Nkx2.5-CRE* hearts at E12.5 and N=3 control and 5 *Phd2^fl/fl^;Nkx2.5-CRE* hearts at E13.5. **E** Quantification of the NOTCH1+16:*LacZ*-positive area on the ventral aspect of the same E12.5 and E13.5 hearts. NS = not significant (unpaired *t* test), error bars show SEM. **F** E15.5 *Phd2^fl/fl^* and *Phd2^fl/fl^;Nkx2.5-CRE* hearts showing NOTCH1+16:*LacZ* activity. N=6 *Phd2^fl/fl^;Nkx2.5-CRE* hearts, all other hearts shown in Supplemental Figure S9. Scale bars = 1mm.

We first investigated the expression of the endocardial-sprout associated, VEGFA/MEF2-driven, angiogenic HLX-3 enhancer (Figure 4B) in *Phd2^fl/fl^;Nkx2.5-Cre* embryonic hearts relative to controls. Expression of the HLX-3:*LacZ* transgene in control *Phd2^fl/fl^* hearts was predominantly restricted to endocardial-derived vessels on the ventral aspect of the heart, in the septum, and in the ventricular free walls, with transient activity also seen at the angiogenic front of the developing SV-derived plexus (Figure 4 and Supplemental Figures 5-6). Conversely, *Phd2^fl/fl^;Nkx2.5-Cre* hearts showed expansion of HLX-3:*LacZ* activity throughout embryonic coronary vessel development. At E11.75, a timepoint in which SV-derived and endocardial-derived EC sprouting first occurs, increased HLX-3:*LacZ* expression was detected in *Phd2^fl/fl^;Nkx2.5-Cre* hearts along the midline on both the ventral and dorsal sides of the heart in a pattern characteristic of endocardial vessel sprouting (Figure 4C and Supplemental Figure 5A). This increased activity was more notable by E12.5, with X-gal expressing endocardial-associated sprouts on the ventral aspect significantly more widespread in *Phd2^fl/fl^;Nkx2.5-Cre* hearts compared to controls (Figure 4D-E and Supplemental Figure 5B). By E13.5 more intense HLX-3:*LacZ* activity was also notable within the SV-derived plexus in the *Phd2^fl/fl^;Nkx2.5-Cre* hearts (Figure 4F). Enhancer activity was also seen in regions not usually populated by either endocardial-derived or SV-derived plexuses, particularly on the ventral aspect and the dorsal apex regions at E12-E13.5 (Figure 4F-G and Supplemental Figure 6). Transverse sections through these hearts demonstrate an increase in HLX-3:*LacZ* activity in the endocardium itself (Figure 4G), and a greater proportion of the sections were populated by HLX-3:*LacZ-*positive vessels, indicating angiogenic expansion of the vascular network (Figure 4H). This widespread activity of the MEF2-driven angiogenic pathway may be a consequence of ectopic activation of endocardial sprouting distal from the normal septum region.

*Phd2^fl/fl^;Nkx2.5-Cre* hearts continued to show increased HLX-3:*LacZ* activity at E15.5 across the entire surface of the ventricles, indicating sustained VEGFA-MEF2 pathway signalling and increased sprouting angiogenesis in the hypoxic heart compared with controls (Figure 4I and Supplemental Figure 7). This was not detected in *Phd2^fl/+^;Nkx2.5-Cre* hearts treated with tamoxifen, suggesting that Cre toxicity is not involved (Supplemental Figure 7A, litter 5). The expanded expression of HLX-3:*LacZ* in *Phd2^fl/fl^;Nkx2.5-Cre* hearts included a greater number of HLX-3:*LacZ*-expressing ECs in the septum region and an increase in HLX-3:*LacZ*-expressing ECs with blood island morphology on the ventral surface, both traditionally associated with endocardial-derived sprouting (Figure 4I and Supplemental Figure S7B). Together, these results indicate that the VEGFA-MEF2 regulatory pathway is over-activated within both the developing endocardial-derived and SV-derived coronary EC populations in response to myocardial hypoxia. However, while this results in expanded ectopic endocardial sprouting, the sustained VEGFA-MEF2 activity in the SV-derived plexus did not appear to lead to expanded SV-derived coronary sprouting.

To better examine the role of hypoxia in SV-derived sprouting, we next investigated the expression of the SOXF-driven Notch1+16 enhancer (Figure 5A) in *Phd2^fl/fl^;Nkx2.5-Cre* embryonic hearts relative to controls. In the heart, this SOXF-driven enhancer becomes active in the SV-derived vascular plexus as it sprouts from the SV at around E11.75 and remains active as this plexus expands to cover the dorsal ventricle at E13.5 before slowly becoming specific to arterial endothelium by birth (Figure 5 and ^13^). This expression closely mimics that of SOX17 (Supplementary Figure 4,^8,36^). As expected, strong Notch1+16:*LacZ* activity was seen in the SV-derived plexus as it sprouts at E12.5-E13.5. However, the expression of the Notch1+16:*LacZ* transgene was not expanded in the SV-derived plexus in *Phd2^fl/fl^;Nkx2.5-Cre* hearts comparative to controls (Figure 5B-D). Although variations in the extent of SV-derived sprouting within and between litters at this stage could potentially mask minor differences, quantification of the area of the dorsal ventricles covered by Notch1+16:*LacZ*-positive vascular plexus at E12.5 and E13.5 found little difference between the hypoxic and control hearts, with a trend for reduced ventricular coverage by the SV-derived plexus (Figure 5D). Notably, unlike for the MEF2-driven HLX-3 enhancer, there was no activation of the Notch1+16 enhancer in the regions beyond the SV-derived plexus on the dorsal ventricular surface, suggesting that the ectopic vessel growth detected here was not utilizing a SOXF-driven transcriptional pathway. Indeed, by E15.5 we detected clear reduced, rather than increased, lateral growth of the SV-derived plexus along the surface of the ventricle in *Phd2^fl/fl^;Nkx2.5-Cre* hearts, with the right ventricle consistently showing less coverage by the SV-derived plexus in *Phd2^fl/fl^;Nkx2.5-Cre* hearts. This is most notable on the ventral aspect, where the SV-derived plexus can be seen extending around the lateral portion of the right ventricle on control *Phd2^fl/fl^*hearts, but was absent or reduced in *Phd2^fl/fl^;Nkx2.5-Cre* hearts (Figure 5F). Overall, these results are in stark contrast to expression of the angiogenic MEF2-driven enhancer at this timepoint, which was strongly increased (alongside increase blood islands) on the lateral portion of the right ventricle in *Phd2^fl/fl^;Nkx2.5-Cre* hearts.

In addition to the SV-derived plexus, in wildtype hearts the SOXF-driven Notch1+16:*LacZ* transgene is also active in a subset of ECs sprouting from the endocardium/septum onto the ventral surface along the septum from E12.5. This correlates with endogenous SOX17, whose expression in this region has previously been associated with “activated” ECs sprouting from the endocardium/septum (Supplemental Figure 4 and ^8^). However, despite the association of hypoxia and VEGFA with increased endocardial-derived sprouting and SOX17-expressing activated ECs, and the increase in angiogenic ECs and MEF2-driven enhancer activity in *Phd2^fl/fl^;Nkx2.5-Cre* hearts, we found no significant qualitative or quantitative differences in Notch1+16:*LacZ* activity in the endocardial-sprout associated ventral vascular regions at E12.5 or E13.5 (Figure 5E). Collectively, these results suggest that stabilized HIFα and subsequent altered VEGFA expression in the heart differentially affected the MEF2-dependent and SOXF-dependent coronary regulatory pathways.

### Arterial maturation is disrupted in *Phd2^fl/fl^;Nkx2.5-Cre* hearts

We next investigated whether the patterning of early arterial development was altered by the increased HIFα stability and altered VEGFA expression in *Phd2^fl/fl^;Nkx2.5-Cre* hearts during development. Coronary ECs derived from both SV and endocardium contribute to the formation of the coronary arteries and veins ^12,14^. Coronary arterial differentiation begins at E12.5-E13.5 as a subset of ECs within both the SV-derived and endocardial-derived plexuses become pre-arterial ECs, reducing cell cycling and inducing the expression of some arterial genes (e.g. *Cxcr4, Efnb2, Dll4*). These pre-arterial ECs then gradually coalesce to form the coronary arteries, with clearly discernible arterial structures seen by E15.5 ^11–13^. Analysis of similar arterial transitions in mouse retinal models have implicated hypoxia/VEGFA-induced angiogenic ECs as an essential precursor for this transition, alongside active Notch1 and high expression of SOX17 and DLL4 ^3,13,37^. Supporting this, our own previous research identified an enhancer for *Dll4* (named Dll4-12), regulated by a combination of VEGFA, SOXF and NOTCH signalling and selectively activated as pre-arterial ECs become specified (Figure 6A)^13,25^.

**Figure 6:**
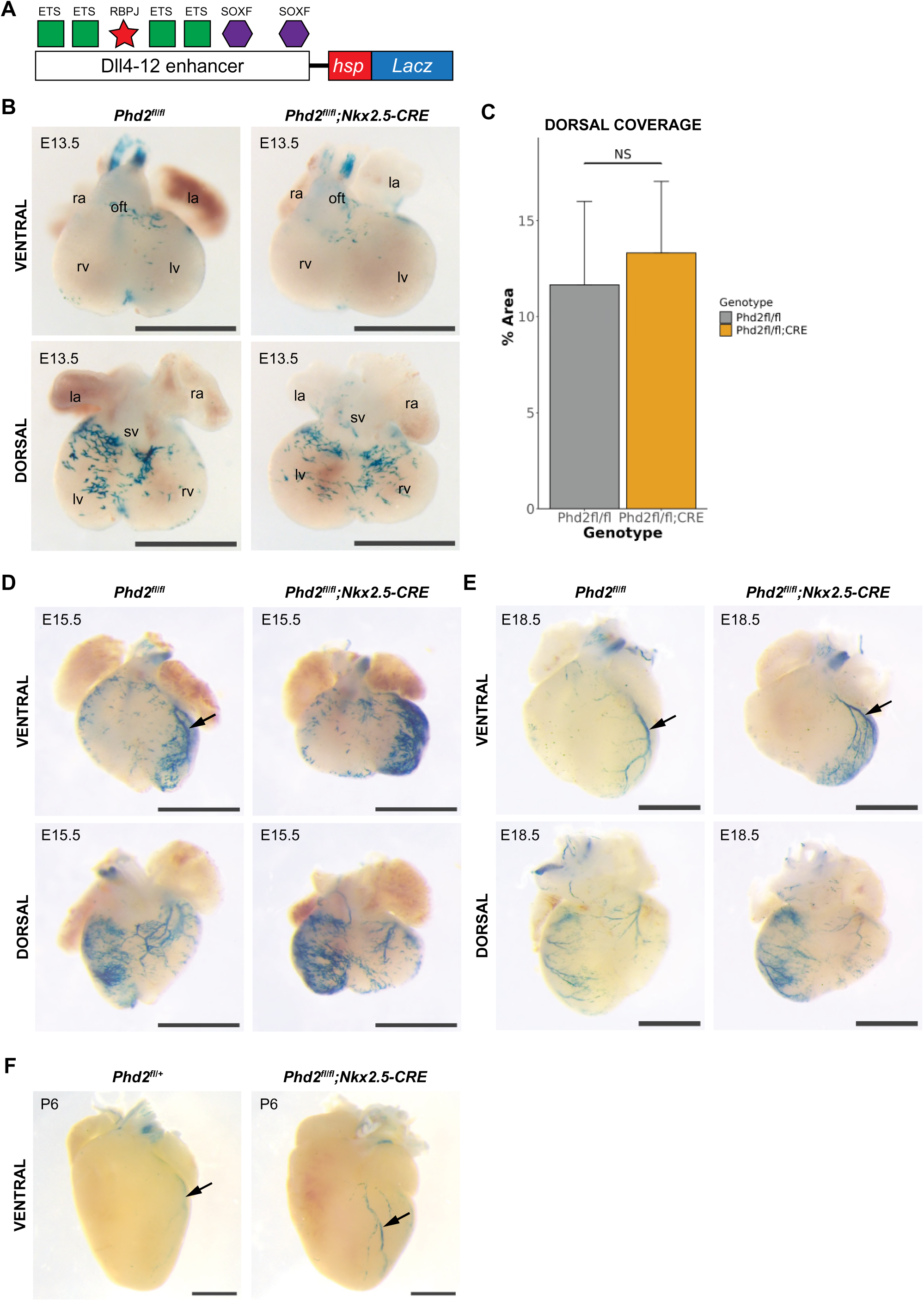
Arterial maturation, but not SOXF-dependent specification, is disrupted in hypoxic hearts **A** Schematic of the Dll4-12:*LacZ* transgene, with the mouse Dll4-12 enhancer sequence cloned upstream of the hsp68 minimal promoter and *LacZ* reporter gene. Validated ETS, SOXF and RBPJ TF binding sites within the enhancer are represented by coloured shapes ^25^. **B** Representative ventral and dorsal views of E13.5 littermate hearts with Dll4-12:*LacZ* activity (blue XGal stain) on *Phd2^fl/fl^* and *Phd2^fl/fl^;Nkx2.5-CRE* genetic backgrounds. N= 4 *Phd2^fl/fl^;Nkx2.5-CRE* hearts, all other hearts shown in Supplemental Figure S10. Scale bars = 1mm. ra = right atrium; la = left atrium; rv = right ventricle; lv = left ventricle; sv = sinus venosus; oft = outflow tract. **C** Quantification of Dll4-12:*LacZ*-positive area on dorsal aspect of the heart, as a percentage of total area. NS = not significant (unpaired *t* test), error bars show SEM, N = 4 *Phd2^fl/fl^;Nkx2.5-CRE* hearts and 5 *Phd2^fl/fl^* hearts. **D** E15.5 *Phd2^fl/fl^*and *Phd2^fl/fl^;Nkx2.5-CRE* hearts showing Dll4-12:*LacZ* activity. N=8 *Phd2^fl/fl^;Nkx2.5-CRE* hearts, all other hearts shown in Supplemental Figure S11, including heterozygous *Phd2^fl/+^;Nkx2.5-CRE* hearts as CRE controls. Scale bars = 1mm. **E** E18.5 *Phd2^fl/fl^*and *Phd2^fl/fl^;Nkx2.5-CRE* hearts showing Dll4-12:*LacZ* activity. N=5 *Phd2^fl/fl^;Nkx2.5-CRE* hearts, all other hearts shown in Supplemental Figure S12A. Scale bars = 1mm. **F** P6 *Phd2^fl/fl^* and *Phd2^fl/fl^;Nkx2.5-CRE* hearts showing Dll4-12:*LacZ* activity. N=5 *Phd2^fl/fl^;Nkx2.5-CRE* hearts collected at P6-8, all other hearts shown in Supplemental Figure S12B.

In wildtype hearts, the Dll4-12:*LacZ* transgene becomes active in pre-arterial ECs at approximately E13.5, remaining specifically expressed in arterial ECs as they migrate and coalesce into mature arterial networks during later embryonic development and into the neonatal stages (Figure 6B and ^13^). Whilst our scRNA-seq detected a slight increase in the ratio of pre-arterial ECs in *Phd2^fl/fl^;Nkx2.5-Cre* hearts at E13.5 (Figure 3E), we did not see significant differences in Dll4-12:*LacZ* transgene activity on the dorsal aspect of *Phd2^fl/fl^;Nkx2.5-Cre* hearts compared with littermate *Phd2^fl/fl^* controls (Figure 6B-C and Supplemental Figure 10). However, variations in Dll4-12:*LacZ* expression between litters at this timepoint mean we may not have detected small differences in expression patterns (Supplemental Figure 10). Notably, from E15.5 we saw significantly increased Dll4-12:*LacZ* activity in *Phd2^fl/fl^;Nkx2.5-Cre* hearts relative to controls, alongside a reduced number of differentiated arterial structures (Figure 6D-F and Supplemental Figure 11-12). For example, the developing left anterior descending artery was consistently under-developed or missing in *Phd2^fl/fl^;Nkx2.5-Cre* hearts when compared with littermate controls, whilst the intensity of Dll4-12:*LacZ* in surrounding ECs was much stronger. Similarly, the dorsal side of the heart had fewer Dll4-12:*LacZ*-positive arterial ECs organising into larger vessels, with greater numbers of Dll4-12:*LacZ*-positive ECs remaining in more discontinuous plexus (Figure 6D and Supplemental Figure 11). Overall, this data suggests that while differentiation of arterial ECs was not overly impacted by stabilized HIF/increased VEGFA levels, the ability of these ECs to migrate and coalesce into mature arteries was compromised. Similar defects were detected at E18.5, where the failure of Dll4-12:*LacZ*-positive ECs to mature and organise into larger coronary arteries in *Phd2^fl/fl^;Nkx2.5-Cre* hearts was even more pronounced (Figure 6E and Supplemental Figure 12). The left anterior descending artery was either narrowed or missing, with increased numbers of small arterioles branching off, again indicating disruption to the process of arterialisation (Figure 6E and Supplemental Figure 12). The coronary arteries located on the dorsal aspect of the heart also appeared narrower and extended a shorter distance towards the apex in comparison to littermate controls. Interesting, these differences did not persist into postnatal life, and hearts collected at postnatal days (P)6 and P8 expressed Dll4-12:*LacZ* in mature coronary arteries in both *Phd2^fl/fl^;Nkx2.5-Cre* and control *Phd2^fl/fl^* hearts (Figure 6F-G and Supplemental Figure 12), indicating that perinatal hearts eventually compensate for the developmental disruption and form a mature vascular network that enables survival.

Overall, our results suggest that early arterial EC differentiation from the plexus is not significantly impacted in a HIFα-stabilized high-VEGFA myocardial environment, despite increased angiogenesis. However, coronary artery maturation, patterning and remodelling are disrupted by this altered microenvironment, resulting in smaller and fewer coronary arteries by the end of gestation, which must be compensated for after birth.

### Venous development is not disrupted in *Phd2^fl/fl^;Nkx2.5-Cre* hearts although patterning is affected

Finally, we investigated whether coronary vein differentiation and maturation was affected in our *Phd2^fl/fl^;Nkx2.5-Cre* hearts. Similar to coronary arterial ECs, coronary vein ECs form from both SV-derived and endocardial-derived ECs via differentiation, although this process occurs slightly later and was therefore not captured by our scRNA-seq analysis. Although the SV ECs from which the dorsal plexus is formed are venous in origin, transcriptomic analysis suggests that these ECs first de-differentiate before re-differentiating into venous coronary ECs ^12^. BMP-SMAD signalling has been implicated in the regulation of vein specification in both systemic and coronary vasculature^13,26^, in part by activating expression of *Coup-TFII* (*Nr2f2*) and *Ephb4*, both key regulators of venous identity^12^. Here, we used the expression of the BMP-SMAD-driven Ephb4-2:*LacZ* transgene line to investigate the effects of hypoxic myocardium on this developmental pathway (Figure 7A)^26,38^. As expected due to its venous origin, the Ephb4-2:*LacZ* transgene is initially strongly expressed in the SV and in the SV-derived plexus, switching off in the plexus around E12.5-E13.0 as the ECs de-differentiate and lose venous identity, although some X-Gal staining remains at E13.5 due to the half-life of β-Gal activity (Figure 7B and ^13^). Here, we found that Ephb4-2:*LacZ* activity was similar in E13.5 *Phd2^fl/fl^;Nkx2.5-Cre* hearts and controls, with X-gal staining largely restricted to similarly migrated SV-derived plexuses in both conditions with no statistical differences in the coverage of the dorsal aspect (Figure 7B-C and Supplemental Figure S13). This supports our observation that SV-derived sprouting was not expanded by alterations to HIFα stability/increased VEGFA expression, and suggests that the ectopic angiogenesis/MEF2 pathway activation seen beyond the plexus did not involve activation of BMP-SMAD signalling. We next looked at E15.5, a timepoint by which definitive venous coronary vessels are formed. At this stage, both *Phd2^fl/fl^;Nkx2.5-Cre* and *Phd2^fl/fl^* control hearts showed selective activity of the Ephb4-2:*LacZ* transgene in the newly formed coronary veins, suggesting that the switch to venous fate driven by BMP-SMAD is not ablated by hypoxic-like conditions in the myocardium. However, the coronary veins in *Phd2^fl/fl^;Nkx2.5-Cre* hearts were truncated and mis-patterned, and the large coronary vein seen on the dorsal aspect of the left ventricle of littermate control hearts was absent in all *Phd2^fl/fl^;Nkx2.5-Cre* hearts (n=7, Figure 7D and Supplemental Figure S13). In addition to expression in the differentiating coronary veins, Ephb4-2:*LacZ* transgene activity was increased on the ventral aspect of both ventricles in all *Phd2^fl/fl^;Nkx2.5-Cre* hearts. This was in a dispersed and punctate pattern that correlates with the location of ectopic/increased VEGFA-MEF2 activity also seen in *Phd2^fl/fl^;Nkx2.5-Cre* (Figure 4I). This suggests that the ectopic sprouting vessels from the endocardium may result in upregulation of the vein associated enhancer in response to hypoxia. Increased Ephb4-2:*LacZ* expression around the pulmonary trunk in a significant subset of *Phd2^fl/fl^;Nkx2.5-Cre* hearts relative to controls also indicates increased formation of small coronary veins in this region (Figure 7 and Supplemental Figure S13). By E18.5, mature EphB4:*LacZ*-expressing coronary veins were seen in both *Phd2^fl/fl^;Nkx2.5-Cre* and control hearts (Figure 7E and Supplemental Figure 14). However, the branching pattern was altered: in *Phd2^fl/fl^* hearts, the large coronary veins extending towards the apex were predominantly located on the left ventricle, whilst in *Phd2^fl/fl^;Nkx2.5-Cre* hearts these veins extended further towards the right ventricle (Figure 7E). However, the ectopic and disparate activity of the enhancer seen on the ventral aspect at E15.5 was largely gone, suggesting this was a transient upregulation of the pathway that does not produce additional coronary veins (Figure 7E and Supplemental Figure 14).

**Figure 7:**
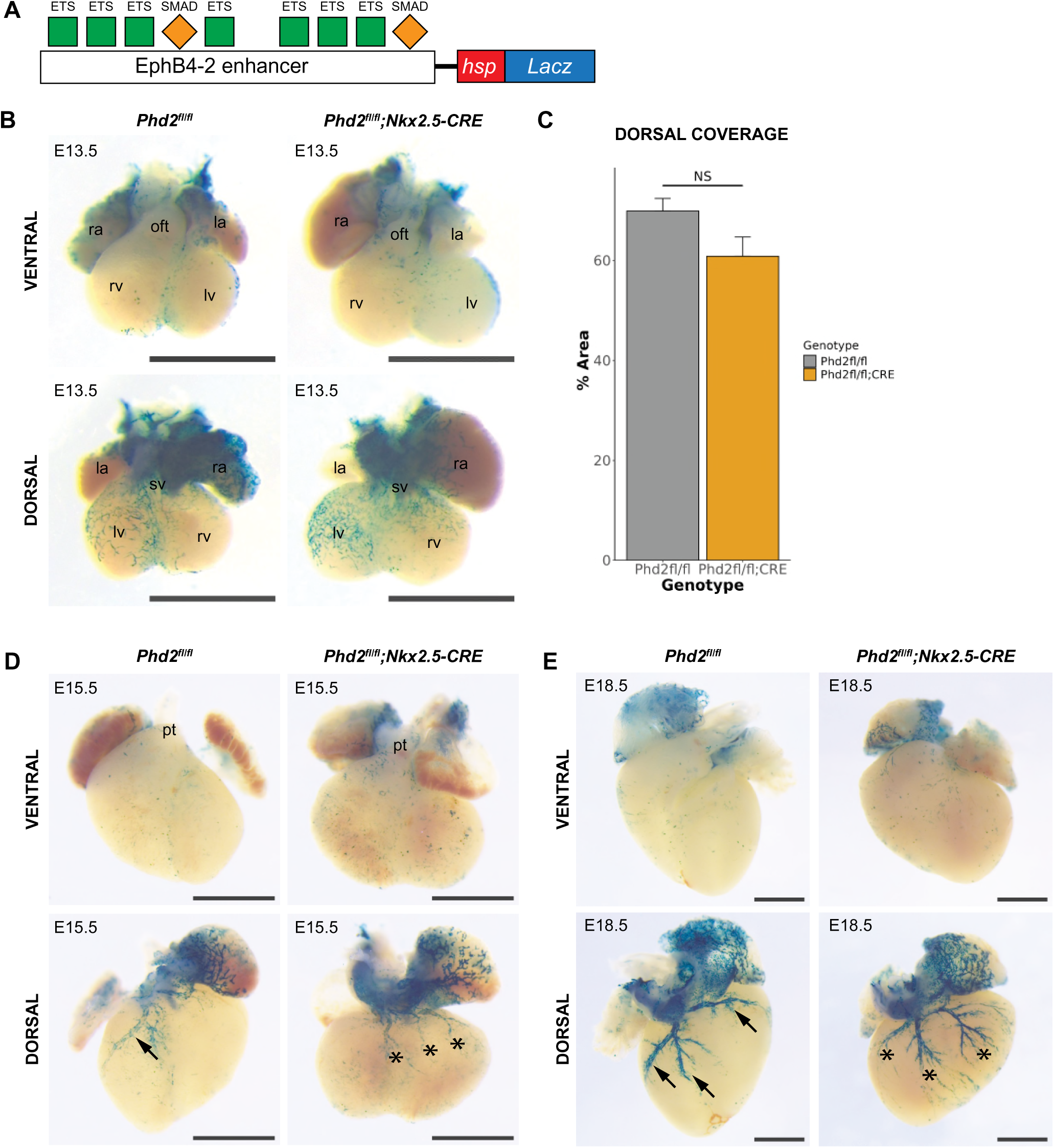
Venous specification and patterning in hypoxic myocardium **A** Schematic of the EphB4-2:*LacZ* transgene, with the mouse EphB4-2 enhancer sequence cloned upstream of the hsp68 minimal promoter and *LacZ* reporter gene. Validated ETS and SMAD TF binding sites within the enhancer are represented by coloured shapes ^26^. **B** Representative E13.5 littermate hearts with EphB4-2:*LacZ* activity (blue XGal stain) on *Phd2^fl/fl^* and *Phd2^fl/fl^;Nkx2.5-CRE* genetic backgrounds. N=6 *Phd2^fl/fl^;Nkx2.5-CRE* hearts, all other hearts shown in Supplemental Figure S13A. Scale bars = 1mm. ra = right atrium; la = left atrium; rv = right ventricle; lv = left ventricle; sv = sinus venosus; oft = outflow tract. **C** Quantification of the extension of the EphB4-2:*LacZ*-positive SV-derived vascular plexus as a percentage of the total dorsal area of *Phd2^fl/fl^* and *Phd2^fl/fl^;Nkx2.5-CRE* hearts at E13.5. NS = not significant (unpaired *t* test), error bars show SEM, N=6 hearts for each genotype. **D** E15.5 *Phd2^fl/fl^* and *Phd2^fl/fl^;Nkx2.5-CRE* hearts showing EphB4:*LacZ* activity – major coronary veins are highlighted with black arrows, with truncated and misdirected veins highlighted with asterisks. N=8 *Phd2^fl/fl^;Nkx2.5-CRE* hearts, all other hearts shown in Supplemental Figure S13B. Scale bars = 1mm. pt = pulmonary trunk. **E** E18.5 *Phd2^fl/fl^* and *Phd2^fl/fl^;Nkx2.5-CRE* hearts showing EphB4-2:*LacZ* activity. N=4 *Phd2^fl/fl^;Nkx2.5-CRE* hearts, all other hearts shown in Supplemental Figure S14. Scale bars = 1mm.

Together, these results indicate that hypoxia does not inhibit the BMP4-SMAD1/5 pathway in driving acquisition of venous identity in coronary ECs, and can transiently drive increased, ectopic activity. However, there is disruption to the patterning of the coronary veins, potentially as a consequence of the thin myocardial wall seen in *Phd2^fl/fl^;Nkx2.5-Cre* hearts.

## DISCUSSION

The link between a hypoxic microenvironment, VEGFA and vascular growth in the systemic vasculature is well established. However, the exact mechanisms by which hypoxia, and the resulting stabilization of HIFα factors, influence coronary vessel growth in the heart have been challenging to ascertain. These are important pathways to understand: hypoxia is an immediate and damaging consequence of myocardial infarction, yet neovascular growth in the ischemic adult heart is insufficient. The research here adds to a growing body of literature linking myocardial hypoxia specifically to endocardial-derived coronary vascularization. Wu et al.,^6^ built on observations of hypoxia-dependent myocardial VEGFA gradients^39^ to demonstrate a direct link between VEGFA-VEGFR2 signalling and endocardial-derived angiogenesis. Supporting this, Sharma et al.^8^ later linked the expansion of endocardial-derived coronary angiogenesis that occurs in response to disrupted SV-derived coronary vasculature growth to increased myocardial hypoxia. Our previous studies of the transcriptional link between VEGF signalling and angiogenic gene expression have also indicated a direct connection between hypoxia and endocardial sprouting, with strong activity of VEGFA-MEF2-driven angiogenic enhancers occurring in regions associated with endocardial-derived sprouting ^13,23^. Here we used a genetic model to mimic hypoxia through ectopic stabilisation of HIFα in the myocardium, enabling us to study the consequences of hypoxia without depleting any of the vascular beds in the developing heart. Our results found increased VEGFA and pro-angiogenic gene expression in these hearts, alongside expanded activity of the VEGFA-MEF2 angiogenic regulatory pathway in a pattern indicative of increased endocardial-derived sprouting. These observations further emphasize a direct and specific link between hypoxia and endocardial coronary vessel sprouting.

It is still not clear how distinct angiogenic sprouting from the endocardium is achieved in either physiological or hypoxic/VEGFA-abundant environments. Both SV-derived and endocardial-derived coronary sprouting occur via angiogenesis by definition, with new vessels forming from existing ones. Whilst previous work implicates VEGFC and ELABELA as the main signalling ligands for SV-derived coronary angiogenesis ^40^, the SV-derived plexus expresses many genes associated with VEGFA-driven angiogenesis in other vascular beds (e.g. *Esm1*, *Apln*, *Pdgfb*, *Sox7, Sox17 and Sox18* ^8,9,36,41–43)^. Further, our previous work also found transient activity of the VEGFA-MEF2-driven angiogenic pathway at the leading edge of the SV-derived plexus as it migrated across the dorsal aspect, indicating that SV-derived sprouting may have some VEGFA dependency. However, we found no evidence of increased SV-derived plexus sprouting in our *Phd2^fl/fl^;Nkx2.5-Cre* hearts. Although we saw expanded activity of the VEGFA-MEF2-driven transgene on the dorsal apex and lateral regions, this was not in a pattern indicative of an expanded SV-derived plexus. Instead, cross-sectional analysis demonstrated that this increased activity was centred on the endocardium: in control hearts, activity of the VEGF-MEF2 driven pathway in the endocardium was restricted to the regions around the septum, whereas in *Phd2^fl/fl^;Nkx2.5-Cre* hearts this expression was expanded to much of the endocardium and into related vessel sprouts. This lack of hypoxic-impact on the SV-derived plexus was further emphasized by the activity of the Notch1+16 and Ephb4-2 enhancers, active in the early SV-derived plexus and not expanded in *Phd2^fl/fl^;Nkx2.5-Cre* hearts. Analysis of an alternate genetic model of hypoxia, using *Mef2c-Cre* to delete VHL and stabilize HIFα, observed both increased endocardial-derived angiogenesis alongside a reduced SV-derived plexus ^35^. While they hypothesised this may be due to a premature migration of the SV-derived vessels into the myocardium, this does not match the pattern of expanded VEGFA expression in these hearts and we saw no evidence of this. Together, our data instead suggests that the genetic pathways regulating angiogenesis from the SV are independent from hypoxia and cognate expanded VEGFA.

Our observations closely align with a previous model in which physiological triggers in the heart exclusively affect endocardial-derived sprouting ^8^. However, although earlier research implicated SOX17 as the specific transcriptional regulator of hypoxia-driven endocardial sprouting^8^, SOX17 is expressed both in endocardial-derived sprouting ECs and throughout the SV-derived plexus at these timepoints (^8^ and this paper). It is therefore highly likely that additional transcription factors are involved in this pathway, permitting SOX17 and other SOXF factors to transactivate differential gene expression patterns in response to signals from either the epicardial surface (in SV-derived vessels) or from within the myocardium (in endocardial-derived vessels). This is supported by our observation that the SOXF-dependent Notch1+16 enhancer was not expanded in our *Phd2^fl/fl^;Nkx2.5-Cre* hearts, despite the expanded SOX17 expression seen in the hypoxic endocardial-derived plexus ^8^.

Hypoxia and subsequent increased VEGFA levels have been closely linked with arterial gene expression and differentiation in other vascular beds ^44^. Although we did not detect a clear difference in genes associated with arterio-venous identity in our transcriptomic analysis, there were moderate shifts in the proportions of coronary ECs towards the pre-arterial clusters and away from venous clusters. This correlated with an observed increase in Dll4-12:*LacZ*-positive ECs away from defined vessels in E15.5 and E18.5 *Phd2^fl/fl^;Nkx2.5-Cre* hearts, potentially reflecting increased numbers of pre-arterial or arterial ECs. Although we did not detect obvious differences in Dll4-12:*LacZ*-positive pre-arterial ECs at E13.5, the stage in which they first differentiate, there was considerable variation in expression across litters at this time-point which could have masked small differences. Interestingly, the increase in Dll4-12:*LacZ*-positive ECs in later stage *Phd2^fl/fl^;Nkx2.5-Cre* hearts correlated with a clear reduction in mature coronary arteries. This indicates that while the differentiation of venous ECs into pre-arterial ECs is not significantly influenced by increased hypoxia and subsequently higher VEGFA levels, the ability of arterial-fated ECs to migrate and coalesce into mature arteries is compromised in the hypoxic/stabilized HIFα environment.

The exact mechanism of how and why the SV-derived endothelial plexus does not expand in response to hypoxic stimuli may be crucial to our understanding of the supressed neovascular response following MI. Lineage tracing studies have confirmed angiogenic sprouting from pre-existing coronary ECs makes a significant contribution to the neovascular response ^45^ yet our previous work found that the VEGFA-MEF2 pathway was not active in the blood vessels surrounding the infarction in a mouse model of MI ^13^. It has also been reported that whilst coronary ECs transiently activate Apelin in response to VEGF in the adult heart, they fail to proliferate and give rise to new vessels, highlighting a reduced angiogenic potential of the myocardium compared with skeletal muscle ^18^. The underlying cause of this difference has not been elucidated, but it has been proposed to be a protective mechanism against tumour angiogenesis within the heart. The lack of MEF2-driven angiogenic gene expression has also been associated with potential HDAC-mediated inhibition of MEF2 activity: increased HDAC activity has been reported in the murine heart post-MI ^46^, and anti-HDAC treatment after MI results in improvements in neovascularisation and cardiac function ^47^. A better understanding of why SV-derived vessels are not sensitive to hypoxia, and why the hypoxia-responsive VEGFA-MEF2 angiogenic pathway is not activated in the ischemic adult heart, would therefore be hugely beneficial in the search for therapeutic interventions to improve neovascularisation post-MI.

## Supporting information

All Supplemental Data

## Non-standard Abbreviations and Acronyms

EC: Endothelial cell
SV: Sinus venosus
E: Embryonic day

## ACKNOWLEDGEMENTS

We would like to extend our sincere thanks to Peter Ratcliffe for providing the *Phd2^flox/flox^* mice, to Shankar Srinivas for providing the *Nkx2.5-Cre* mice and Xin Lu for providing the Rosa26-*LacZ* reporter mice.

## SOURCES OF FUNDING

This work was funded by grants from the British Heart Foundation (FS/17/35/32929 and FS/SBSRF/22/31037 for S.D.; FS/18/62/33967 for S.P; PG/21/10704 for D.S), the BHF Centre of Research Excellence Oxford (RE/13/1/30181) and the Fondation Leducq (S.B.).

## DISCLOSURES

None.

## SUPPLEMENTAL MATERIAL

Tables S1–S3

Figures S1-S14

## NOVELTY AND SIGNIFICANCE

### What is known?

- Hypoxia is an immediate and damaging consequence of myocardial infarction.
- The neo-vascular response in the ischaemic adult heart is insufficient, impeding the ability of the heart to repair itself following myocardial infarction.
- Hypoxia, and resultant increased VEGFA expression, can stimulate angiogenesis in the systemic vasculature, but their influence on different aspects of coronary vessel has been more difficult to assess.

### What new information does this article contribute?

- Hypoxic myocardium in the developing heart increased pro-angiogenic gene expression and shifted endothelial cells towards a non-proliferative tip-cell phenotype.
- A VEGFA-MEF2 angiogenic regulatory pathway expanded in hypoxic hearts in a pattern indicative of increased endocardial-derived sprouting, whilst SV-derived vessel growth was unaffected.
- Arterio-venous differentiation was unaffected by hypoxic myocardium, but later large vessel maturation and patterning was disrupted.

### Summary of Novelty and Significance

This paper provides new insights into the mechanisms linking different types of coronary vascular growth and hypoxia, with relevance to research topics spanning classical developmental biology, transcriptional regulation of vascular growth, cardiovascular responses to ischemic injury and regenerative medicine.

We used a genetic model to mimic myocardial hypoxia through ectopic stabilisation of HIFα, enabling us to study the consequences of hypoxia on different aspects of coronary vessel formation. This analysis combined single cell transcriptomics with an examination of the activity of multiple enhancer:reporter transgenes in experimental hearts relative to littermate controls. This approach is especially powerful in the heart, as it enables us to directly examine the consequences of hypoxia on different coronary vascular beds forming from distinct origins via disparate signalling and transcriptional pathways. Our results found increased pro-angiogenic gene expression in these hearts, alongside expanded activity of the VEGFA-MEF2 angiogenic regulatory pathway in a pattern indicative of increased endocardial-derived sprouting. Conversely, despite the expanded VEGFA expression, angiogenesis from the sinus venosus-derived vascular plexus was relatively unaffected in these hypoxic conditions. Initial arteriovenous specification also occurred normally, but mature coronary artery formation was delayed, as the ability of arterial-fated ECs to migrate and coalesce into mature arteries was compromised in the hypoxic myocardium.

## Notes

### Competing Interest Statement

The authors have declared no competing interest.

