## Supplementary material for "Hypoxia differentially affects coronary vessel formation during heart development": All Supplemental Data

**Supplemental Table S1: Differentially expressed genes across all cells**

Differential gene expression analysis results comparing all *Phd2*$^{fl/fl}$*;Nkx2.5-Cre* with control *Phd2*$^{fl/fl}$ cells, showing 37 genes upregulated in *Phd2*$^{fl/fl}$*;Nkx2.5-Cre* cells and 8 genes downregulated in *Phd2*$^{fl/fl}$*;Nkx2.5-Cre* cells. Genes were included if adjusted p-value after multiple testing correction < 0.05 and log2 fold change > 0.2.

| Gene | p_val | avg_log2FC | pct.1 | pct.2 | p_val_adj | Direction |
|---|---|---|---|---|---|---|
| ***Tnnt2*** | 5.83E-07 | 0.769966 | 0.245 | 0.197 | 0.018815 | Upregulated |
| ***Pgk1*** | 2.75E-52 | 0.564618 | 0.816 | 0.718 | 8.86E-48 | Upregulated |
| ***Mif*** | 4.77E-34 | 0.527886 | 0.834 | 0.754 | 1.54E-29 | Upregulated |
| ***Aldoa*** | 1.16E-50 | 0.520576 | 0.846 | 0.745 | 3.74E-46 | Upregulated |
| ***Tpi1*** | 5.19E-42 | 0.502359 | 0.783 | 0.673 | 1.68E-37 | Upregulated |
| ***Ldha*** | 5.24E-39 | 0.490589 | 0.901 | 0.842 | 1.69E-34 | Upregulated |
| ***Ptn*** | 1.71E-11 | 0.46801 | 0.402 | 0.333 | 5.52E-07 | Upregulated |
| ***Bnip3*** | 8.76E-83 | 0.458984 | 0.295 | 0.12 | 2.83E-78 | Upregulated |
| ***Gapdh*** | 1.08E-21 | 0.430632 | 0.937 | 0.918 | 3.47E-17 | Upregulated |
| ***Slc8a1*** | 1.74E-11 | 0.425151 | 0.152 | 0.101 | 5.62E-07 | Upregulated |
| ***Nexn*** | 1.19E-09 | 0.394475 | 0.194 | 0.14 | 3.85E-05 | Upregulated |
| ***Sorbs2*** | 5.45E-18 | 0.385761 | 0.199 | 0.125 | 1.76E-13 | Upregulated |
| ***Sparcl1*** | 1.87E-35 | 0.380434 | 0.342 | 0.209 | 6.03E-31 | Upregulated |
| ***Pkm*** | 9.01E-36 | 0.37744 | 0.807 | 0.706 | 2.91E-31 | Upregulated |
| ***Cryab*** | 1.45E-06 | 0.33295 | 0.146 | 0.109 | 0.04686 | Upregulated |
| ***Vcan*** | 1.61E-08 | 0.315993 | 0.491 | 0.418 | 5.19E-04 | Upregulated |
| ***Pgam1*** | 2.39E-23 | 0.311196 | 0.787 | 0.701 | 7.70E-19 | Upregulated |
| ***Cfh*** | 2.23E-11 | 0.299935 | 0.66 | 0.619 | 7.20E-07 | Upregulated |
| ***Igfbp3*** | 2.73E-10 | 0.298002 | 0.25 | 0.188 | 8.83E-06 | Upregulated |
| ***Lbh*** | 8.02E-13 | 0.279292 | 0.384 | 0.298 | 2.59E-08 | Upregulated |
| ***Slc16a3*** | 5.16E-65 | 0.274954 | 0.249 | 0.101 | 1.67E-60 | Upregulated |
| ***Pfkl*** | 4.29E-55 | 0.259505 | 0.534 | 0.347 | 1.39E-50 | Upregulated |
| ***Slc2a1*** | 1.71E-39 | 0.253781 | 0.294 | 0.165 | 5.52E-35 | Upregulated |
| ***Gpi1*** | 3.69E-27 | 0.244252 | 0.782 | 0.669 | 1.19E-22 | Upregulated |
| ***Dst*** | 7.53E-37 | 0.243859 | 0.773 | 0.636 | 2.43E-32 | Upregulated |
| ***Palld*** | 1.17E-21 | 0.233799 | 0.372 | 0.265 | 3.78E-17 | Upregulated |
| ***Glo1*** | 1.17E-50 | 0.232038 | 0.702 | 0.535 | 3.78E-46 | Upregulated |
| ***Fam162a*** | 6.65E-10 | 0.223962 | 0.777 | 0.701 | 2.15E-05 | Upregulated |
| ***Nrg1*** | 2.36E-10 | 0.219786 | 0.391 | 0.319 | 7.60E-06 | Upregulated |
| ***Cdh2*** | 2.22E-24 | 0.216845 | 0.194 | 0.108 | 7.17E-20 | Upregulated |
| ***Dsp*** | 2.45E-35 | 0.21093 | 0.267 | 0.146 | 7.91E-31 | Upregulated |
| ***Ldhb*** | 9.33E-18 | 0.210919 | 0.357 | 0.256 | 3.01E-13 | Upregulated |
| ***Ddx3y*** | 6.37E-25 | 0.210709 | 0.291 | 0.19 | 2.06E-20 | Upregulated |
| ***Gyg*** | 1.17E-13 | 0.209222 | 0.462 | 0.359 | 3.78E-09 | Upregulated |
| ***Atp2a2*** | 1.24E-10 | 0.206864 | 0.791 | 0.704 | 4.01E-06 | Upregulated |
| ***Tm4sf1*** | 1.61E-10 | 0.202622 | 0.83 | 0.807 | 5.20E-06 | Upregulated |
| ***Bgn*** | 2.77E-12 | 0.201426 | 0.677 | 0.612 | 8.93E-08 | Upregulated |
| ***Rps2*** | 2.46E-131 | -0.43798 | 0.968 | 0.971 | 7.93E-127 | Downregulated |
| ***Hspa1b*** | 2.14E-15 | -0.40377 | 0.495 | 0.559 | 6.90E-11 | Downregulated |
| ***Hspa1a*** | 1.28E-12 | -0.37829 | 0.489 | 0.55 | 4.12E-08 | Downregulated |
| ***Xist*** | 7.10E-26 | -0.30238 | 0.549 | 0.689 | 2.29E-21 | Downregulated |
| ***Hsph1*** | 8.11E-09 | -0.30149 | 0.717 | 0.717 | 2.62E-04 | Downregulated |
| ***Jun*** | 1.95E-11 | -0.26893 | 0.925 | 0.931 | 6.30E-07 | Downregulated |
| ***Igfbp5*** | 5.19E-07 | -0.24301 | 0.65 | 0.685 | 0.01674 | Downregulated |
| ***Gm1673*** | 1.40E-16 | -0.20656 | 0.736 | 0.761 | 4.53E-12 | Downregulated |

**Supplemental Table S2: Pathway analysis of genes upregulated in *Phd2*$^{fl/fl}$*;Nkx2.5-Cre* hearts**

Gene-set enrichment analysis results from pathway analysis using g:Profiler on genes upregulated in hypoxic conditions at E13.5.

| term_id | term_name | p_value |
|---|---|---|
| **GO:0006090** | pyruvate metabolic process | 1.21E-08 |
| **GO:0019674** | NAD metabolic process | 2.30E-08 |
| **GO:0006007** | glucose catabolic process | 2.62E-08 |
| **GO:0019320** | hexose catabolic process | 1.78E-07 |
| **GO:0046365** | monosaccharide catabolic process | 2.46E-07 |
| **GO:0019677** | NAD catabolic process | 4.62E-07 |
| **GO:0061718** | glucose catabolic process to pyruvate | 4.62E-07 |
| **GO:0061621** | canonical glycolysis | 4.62E-07 |
| **GO:0006735** | NADH regeneration | 4.62E-07 |
| **GO:0061620** | glycolytic process through glucose-6-phosphate | 1.22E-06 |
| **GO:0061615** | glycolytic process through fructose-6-phosphate | 4.11E-06 |
| **GO:0046496** | nicotinamide nucleotide metabolic process | 4.77E-06 |
| **GO:0019362** | pyridine nucleotide metabolic process | 4.77E-06 |
| **GO:0072524** | pyridine-containing compound metabolic process | 5.26E-06 |
| **GO:0032787** | monocarboxylic acid metabolic process | 2.39E-05 |
| **GO:0006006** | glucose metabolic process | 2.75E-05 |
| **GO:0006734** | NADH metabolic process | 4.38E-05 |
| **GO:0016052** | carbohydrate catabolic process | 6.89E-05 |
| **GO:0006096** | glycolytic process | 9.52E-05 |
| **GO:0019364** | pyridine nucleotide catabolic process | 1.08E-04 |
| **GO:0009137** | purine nucleoside diphosphate catabolic process | 1.23E-04 |
| **GO:0072526** | pyridine-containing compound catabolic process | 1.23E-04 |
| **GO:0046032** | ADP catabolic process | 1.23E-04 |
| **GO:0009181** | purine ribonucleoside diphosphate catabolic process | 1.23E-04 |
| **GO:0019318** | hexose metabolic process | 1.52E-04 |
| **GO:0009191** | ribonucleoside diphosphate catabolic process | 1.79E-04 |
| **GO:0009134** | nucleoside diphosphate catabolic process | 2.01E-04 |
| **GO:0046031** | ADP metabolic process | 2.01E-04 |
| **GO:0009135** | purine nucleoside diphosphate metabolic process | 2.26E-04 |
| **GO:0009179** | purine ribonucleoside diphosphate metabolic process | 2.26E-04 |
| **GO:0005996** | monosaccharide metabolic process | 2.66E-04 |
| **GO:0009185** | ribonucleoside diphosphate metabolic process | 6.58E-04 |
| **GO:0009132** | nucleoside diphosphate metabolic process | 7.99E-04 |
| **GO:0044282** | small molecule catabolic process | 9.56E-04 |
| **GO:0009154** | purine ribonucleotide catabolic process | 9.65E-04 |
| **GO:0006195** | purine nucleotide catabolic process | 0.001159204 |
| **GO:0072523** | purine-containing compound catabolic process | 0.001268211 |
| **GO:0046166** | glyceraldehyde-3-phosphate biosynthetic process | 0.001510487 |
| **GO:0009261** | ribonucleotide catabolic process | 0.001512467 |
| **GO:0019752** | carboxylic acid metabolic process | 0.0018921 |
| **GO:0043436** | oxoacid metabolic process | 0.002109656 |
| **GO:0006082** | organic acid metabolic process | 0.002287316 |
| **GO:0009166** | nucleotide catabolic process | 0.002928081 |
| **GO:1901292** | nucleoside phosphate catabolic process | 0.00397931 |
| **GO:0005975** | carbohydrate metabolic process | 0.005614229 |
| **GO:0019682** | glyceraldehyde-3-phosphate metabolic process | 0.005882122 |

| | | |
|---|---|---|
| **GO:0046034** | ATP metabolic process | 0.00846916 |
| **GO:0006091** | generation of precursor metabolites and energy | 0.012531191 |
| **GO:0006163** | purine nucleotide metabolic process | 0.01643671 |
| **GO:0046434** | organophosphate catabolic process | 0.019425481 |
| **GO:0009205** | purine ribonucleoside triphosphate metabolic process | 0.020521511 |
| **GO:0046184** | aldehyde biosynthetic process | 0.021375216 |
| **GO:0072521** | purine-containing compound metabolic process | 0.023290652 |
| **GO:0009144** | purine nucleoside triphosphate metabolic process | 0.025458806 |
| **GO:0009199** | ribonucleoside triphosphate metabolic process | 0.026559727 |
| **GO:0006081** | cellular aldehyde metabolic process | 0.029980448 |
| **GO:1901136** | carbohydrate derivative catabolic process | 0.039837804 |
| **GO:0009141** | nucleoside triphosphate metabolic process | 0.043308838 |
| **GO:0009117** | nucleotide metabolic process | 0.044876849 |
| **KEGG:00010** | Glycolysis / Gluconeogenesis | 3.13E-09 |
| **KEGG:05230** | Central carbon metabolism in cancer | 3.28E-04 |
| **KEGG:01230** | Biosynthesis of amino acids | 0.007055504 |
| **KEGG:01200** | Carbon metabolism | 0.008775507 |

**Supplemental Table S3: Top 10 up- and downregulated genes for each cluster of coronary ECs**

Differentially expressed genes for each cluster were ranked by lowest p value or highest log2 fold change, and the top 10 up-and down-regulated genes from each list (with a value of >0.1 in both pct.1 and pct.2) are shown in these tables.

### Cluster 0 – Capillaries (G1)

#### Ranked by p value

| Gene | p_val | avg_log2FC | p_val_adj |
|---|---|---|---|
| ***Gde1*** | 0.00011723 | 0.390050615 | 1 |
| ***Leng1*** | 0.00027847 | 0.272388824 | 1 |
| ***Pdpk1*** | 0.00031951 | 0.336365428 | 1 |
| ***Nt5dc3*** | 0.00036651 | 0.235225613 | 1 |
| ***Kitl*** | 0.00037917 | 0.653991355 | 1 |
| ***Rbbp6*** | 0.00050378 | 0.505719659 | 1 |
| ***Wbp1l*** | 0.00054522 | 0.298302515 | 1 |
| ***S100a1*** | 0.00065138 | 0.293045418 | 1 |
| ***Ypel5*** | 0.00098801 | 0.271896437 | 1 |
| ***Zfp655*** | 0.00119233 | 0.231546291 | 1 |
| ***Rps2*** | 8.57E-08 | -0.49751659 | 0.002767 |
| ***Ube2n*** | 9.28E-06 | -0.470006317 | 0.299742 |
| ***Ipo11*** | 0.00018916 | -0.490648208 | 1 |
| ***Ccdc102a*** | 0.00072934 | -0.323914943 | 1 |
| ***Faf1*** | 0.00128615 | -0.277238636 | 1 |
| ***Cep83*** | 0.0017621 | -0.391576322 | 1 |
| ***Anapc1*** | 0.00185757 | -0.319123911 | 1 |
| ***Ciao2b*** | 0.00226197 | -0.342031002 | 1 |
| ***Rhot1*** | 0.0026321 | -0.269202788 | 1 |
| ***Ciao2b*** | 0.00226197 | -0.342031002 | 1 |

#### Ranked by Log2FC

| Gene | p_val | avg_log2FC | p_val_adj |
|---|---|---|---|
| ***Klf2*** | 0.004689616 | 0.725003885 | 1 |
| ***Kitl*** | 0.000379165 | 0.653991355 | 1 |
| ***Meg3*** | 0.124498511 | 0.566063475 | 1 |
| ***Hba-a1*** | 0.287058091 | 0.538321035 | 1 |
| ***Gm42418*** | 0.019268232 | 0.510193699 | 1 |
| ***Rbbp6*** | 0.000503776 | 0.505719659 | 1 |
| ***Arhgap21*** | 0.001802563 | 0.435348407 | 1 |
| ***Hist1h1b*** | 0.974159252 | 0.434916695 | 1 |
| ***Ddx3y*** | 0.002848712 | 0.433708277 | 1 |
| ***AY036118*** | 0.055592311 | 0.431335085 | 1 |
| ***Hbb-y*** | 0.567172051 | -1.853661647 | 1 |
| ***Xist*** | 0.003345007 | -0.8270122 | 1 |
| ***Cxcl1*** | 0.062312016 | -0.591125334 | 1 |
| ***Rps2*** | 8.57E-08 | -0.49751659 | 0.002766635 |
| ***Ipo11*** | 0.000189157 | -0.490648208 | 1 |
| ***Ube2n*** | 9.28E-06 | -0.470006317 | 0.299741522 |
| ***D10Wsu102e*** | 0.151085311 | -0.464868495 | 1 |
| ***Igfbp3*** | 0.831007544 | -0.409757013 | 1 |
| ***Phlda1*** | 0.067020612 | -0.404353045 | 1 |
| ***Cep83*** | 0.001762101 | -0.391576322 | 1 |

### Cluster 1 – Cycling capillaries (G2M) I

#### Ranked by p value

| Gene | p_val | avg_log2FC | p_val_adj |
|---|---|---|---|
| ***Sugp1*** | 6.20E-05 | 0.24203132 | 1 |
| ***Txnip*** | 0.000278716 | 0.3862888 | 1 |
| ***Igf2*** | 0.000303273 | 0.77567039 | 1 |
| ***Sbds*** | 0.000423675 | 0.349017827 | 1 |
| ***Dstyk*** | 0.000480631 | 0.190215073 | 1 |
| ***Uggt2*** | 0.000550018 | 0.166639742 | 1 |
| ***Irx5*** | 0.000695091 | 0.218097122 | 1 |
| ***Kif1c*** | 0.000874866 | 0.221223692 | 1 |
| ***Socs5*** | 0.000934012 | 0.179392902 | 1 |
| **H19** | 0.001052546 | 0.74331054 | 1 |
| ***Rps2*** | 4.68E-11 | -0.646213199 | 1.51E-06 |
| ***Dynll1*** | 0.000145714 | -0.331629136 | 1 |
| ***Mtf2*** | 0.000256111 | -0.305828937 | 1 |
| ***Snrpd3*** | 0.000295245 | -0.361624668 | 1 |
| ***Tmem115*** | 0.000376663 | -0.193695617 | 1 |
| ***Hmgn5*** | 0.000432631 | -0.439857748 | 1 |
| ***Atp5f1*** | 0.000679567 | -0.309225893 | 1 |
| **Rpl41** | 0.000734855 | -0.188007911 | 1 |
| **Tax1bp3** | 0.001223843 | -0.329453953 | 1 |
| **Rps25** | 0.001255246 | -0.180352122 | 1 |

#### Ranked by Log2FC

| Gene | p_val | avg_log2FC | p_val_adj |
|---|---|---|---|
| ***Cdkn1c*** | 0.005889196 | 1.049048185 | 1 |
| ***Igf2*** | 0.000303273 | 0.77567039 | 1 |
| ***H19*** | 0.001052546 | 0.74331054 | 1 |
| ***Igfbp5*** | 0.012441949 | 0.740472239 | 1 |
| ***Gm42418*** | 0.006286325 | 0.700558665 | 1 |
| ***Hspa1b*** | 0.35804158 | 0.662159348 | 1 |
| ***AY036118*** | 0.356136325 | 0.634708372 | 1 |
| ***Igfbp3*** | 0.377852289 | 0.578337023 | 1 |
| ***Nts*** | 0.489354823 | 0.557281456 | 1 |
| ***Bmp2*** | 0.028958109 | 0.516571197 | 1 |
| ***Fabp4*** | 0.004033197 | -0.753906434 | 1 |
| ***Rps2*** | 4.68E-11 | -0.646213199 | 1.51E-06 |
| ***Hmgn5*** | 0.000432631 | -0.439857748 | 1 |
| ***Khdrbs3*** | 0.062533871 | -0.404020934 | 1 |
| ***Actn1*** | 0.002926682 | -0.378688117 | 1 |
| ***Tma7*** | 0.001730115 | -0.37544394 | 1 |
| ***Meox2*** | 0.039765899 | -0.365470792 | 1 |
| ***Snrpd3*** | 0.000295245 | -0.361624668 | 1 |
| ***Cdr2*** | 0.002326507 | -0.357215652 | 1 |
| ***Myl12a*** | 0.00153072 | -0.353867451 | 1 |

### Cluster 2 – Cycling capillaries (G2M) II

#### Ranked by p value

| Gene | p_val | avg_log2FC | p_val_adj |
|---|---|---|---|
| ***Plvap*** | 0.000205 | 0.388661 | 1 |
| ***Mvb12b*** | 0.000425 | 0.235572 | 1 |
| ***Prpf3*** | 0.000465 | 0.28021 | 1 |
| ***Map3k1*** | 0.000636 | 0.376033 | 1 |
| ***Nol9*** | 0.000706 | 0.323818 | 1 |
| ***Lmf2*** | 0.000782 | 0.301696 | 1 |
| ***Celf1*** | 0.001375 | 0.478418 | 1 |
| ***Cdc42ep1*** | 0.001447 | 0.378949 | 1 |
| ***Nelfb*** | 0.00145 | 0.254816 | 1 |
| ***Inpp5a*** | 0.001526 | 0.24337 | 1 |
| ***Rps2*** | 7.18E-08 | -0.64573 | 0.002318 |
| ***Rps25*** | 0.001115 | -0.23947 | 1 |
| ***Hint2*** | 0.002375 | -0.33962 | 1 |
| ***Rpl41*** | 0.002616 | -0.18317 | 1 |
| ***Peg3*** | 0.002851 | -0.87483 | 1 |
| ***Rnf220*** | 0.003174 | -0.2932 | 1 |
| ***Rplp1*** | 0.005094 | -0.16596 | 1 |
| ***Adamts1*** | 0.006038 | -0.5465 | 1 |
| ***Pck2*** | 0.00612 | -0.28738 | 1 |
| ***Eif2b1*** | 0.0066 | -0.24067 | 1 |

#### Ranked by Log2FC

| Gene | p_val | avg_log2FC | p_val_adj |
|---|---|---|---|
| ***Cdkn1c*** | 0.218081 | 0.978784202 | 1 |
| ***Igf2*** | 0.016468 | 0.870311067 | 1 |
| ***Gpihbp1*** | 0.003234 | 0.851538423 | 1 |
| ***H19*** | 0.009149 | 0.788964926 | 1 |
| ***Mdk*** | 0.160978 | 0.623917553 | 1 |
| ***Apoe*** | 0.234516 | 0.523936274 | 1 |
| ***Ccnd1*** | 0.069655 | 0.500791803 | 1 |
| ***Bst2*** | 0.043076 | 0.481641551 | 1 |
| ***Celf1*** | 0.001375 | 0.478418309 | 1 |
| ***Siva1*** | 0.022822 | 0.45129336 | 1 |
| ***Hspa1a*** | 0.265946 | -0.957860989 | 1 |
| ***Peg3*** | 0.002851 | -0.874834702 | 1 |
| ***Lgals1*** | 0.013413 | -0.822280109 | 1 |
| ***Rps2*** | 7.18E-08 | -0.645726595 | 0.002318 |
| ***Hspb1*** | 0.535309 | -0.624419269 | 1 |
| ***Hist1h3c*** | 0.0729 | -0.611836473 | 1 |
| ***Casp3*** | 0.011168 | -0.551256991 | 1 |
| ***Adamts1*** | 0.006038 | -0.546498283 | 1 |
| ***Hspa1b*** | 0.067881 | -0.533781183 | 1 |
| ***Gadd45g*** | 0.007104 | -0.516875375 | 1 |

### Cluster 3 – Cycling capillaries (S)

#### Ranked by p value

| Gene | p_val | avg_log2FC | p_val_adj |
|---|---|---|---|
| ***Arhgap42*** | 3.94E-05 | 0.41966577 | 1 |
| ***Fbxl3*** | 0.000203 | 0.316617736 | 1 |
| ***Nom1*** | 0.000252 | 0.495964242 | 1 |
| ***Dgkh*** | 0.000411 | 0.502505782 | 1 |
| ***A430106G13Rik*** | 0.000415 | 0.126505997 | 1 |
| ***Ptgr2*** | 0.00046 | 0.219378439 | 1 |
| ***Gpsm3*** | 0.000521 | 0.279574407 | 1 |
| ***A630072M18Rik*** | 0.000745 | 0.212660187 | 1 |
| ***Hipk2*** | 0.000785 | 0.358344944 | 1 |
| ***Mapk6*** | 0.000789 | 0.352950167 | 1 |
| ***Rps2*** | 1.65E-07 | -0.547120526 | 0.00534 |
| ***Rplp2*** | 0.000263 | -0.25675798 | 1 |
| ***Rpl36a*** | 0.000571 | -0.384246547 | 1 |
| ***Rps26*** | 0.001141 | -0.258849648 | 1 |
| ***Rpl31*** | 0.001426 | -0.276450886 | 1 |
| ***Rpl36*** | 0.001426 | -0.353627745 | 1 |
| ***Uqcr10*** | 0.002022 | -0.421236534 | 1 |
| ***Slc25a4*** | 0.002398 | -0.318748639 | 1 |
| ***Rpl38*** | 0.002838 | -0.353345982 | 1 |
| ***Mcm6*** | 0.003769 | -0.486236239 | 1 |

#### Ranked by Log2FC

| Gene | p_val | avg_log2FC | p_val_adj |
|---|---|---|---|
| ***Hist1h1b*** | 0.886983 | 0.722487383 | 1 |
| ***Igfbp3*** | 0.085272 | 0.705670361 | 1 |
| ***Prnd*** | 0.005544 | 0.655083323 | 1 |
| ***Sat1*** | 0.017064 | 0.633576131 | 1 |
| ***Filip1l*** | 0.010444 | 0.603631256 | 1 |
| ***Neurl3*** | 0.076208 | 0.602881781 | 1 |
| ***Hist1h1e*** | 0.047593 | 0.572414219 | 1 |
| ***Igf2*** | 0.095273 | 0.556477231 | 1 |
| ***Hspa1a*** | 0.780404 | 0.527924532 | 1 |
| ***Itm2a*** | 0.064762 | 0.51367167 | 1 |
| ***Lgals1*** | 0.204271 | -0.551189662 | 1 |
| ***Rps2*** | 1.65E-07 | -0.547120526 | 0.00534 |
| ***Edn1*** | 0.314449 | -0.54360743 | 1 |
| ***Top2a*** | 0.289974 | -0.536447741 | 1 |
| ***Ube2c*** | 0.183903 | -0.529443437 | 1 |
| ***Junb*** | 0.055708 | -0.504691134 | 1 |
| ***Mcm6*** | 0.003769 | -0.486236239 | 1 |
| ***Snhg6*** | 0.020293 | -0.483629789 | 1 |
| ***Bok*** | 0.027197 | -0.466545574 | 1 |
| ***Ppih*** | 0.037297 | -0.462537922 | 1 |

### Cluster 4 – Pre-arterial ECs

#### Ranked by p value

| Gene | p_val | avg_log2FC | p_val_adj |
|---|---|---|---|
| *Eif3m* | 0.000119 | 0.554738775 | 1 |
| *Tmem259* | 0.000188 | 0.326529927 | 1 |
| *Nhsl1* | 0.000245 | 0.414126956 | 1 |
| *Washc1* | 0.000431 | 0.327404959 | 1 |
| *Cnot2* | 0.000754 | 0.427832238 | 1 |
| *Sertad2* | 0.00077 | 0.3228216 | 1 |
| *D1Ertd622e* | 0.000778 | 0.527539427 | 1 |
| *Akr1a1* | 0.000785 | 0.362995178 | 1 |
| *Ppp1cb* | 0.000787 | 0.432257916 | 1 |
| *Gm4258* | 0.00096 | 0.490973903 | 1 |
| *Lamb1* | 0.000233 | -0.64519466 | 1 |
| *Plat* | 0.000637 | -0.644919333 | 1 |
| *Rhoc* | 0.000868 | -0.517840713 | 1 |
| *Fam162a* | 0.001309 | -0.464373887 | 1 |
| *Snhg6* | 0.002611 | -0.476571392 | 1 |
| *Ppwd1* | 0.003108 | -0.290945868 | 1 |
| *Nes* | 0.003946 | -0.446447793 | 1 |
| *Sema3f* | 0.004284 | -0.306099168 | 1 |
| *Dnajc24* | 0.004293 | -0.384834972 | 1 |
| *Trib1* | 0.004855 | -0.731839602 | 1 |

#### Ranked by Log2FC

| Gene | p_val | avg_log2FC | p_val_adj |
|---|---|---|---|
| *Tiparp* | 0.009744 | 0.86783055 | 1 |
| *Cavin2* | 0.015429 | 0.606246872 | 1 |
| *Btbd3* | 0.023565 | 0.561620114 | 1 |
| *Eif3m* | 0.000119 | 0.554738775 | 1 |
| *Cdc42ep3* | 0.073479 | 0.54450207 | 1 |
| *Clec14a* | 0.00328 | 0.5434638 | 1 |
| *Cldn5* | 0.016727 | 0.533343382 | 1 |
| *Hist1h2bc* | 0.190422 | 0.528619448 | 1 |
| *D1Ertd622e* | 0.000778 | 0.527539427 | 1 |
| *Abcg2* | 0.014897 | 0.520565794 | 1 |
| *Gm26917* | 0.12925 | -3.095267775 | 1 |
| *Nts* | 0.014544 | -1.884371705 | 1 |
| *AY036118* | 0.010038 | -1.020831048 | 1 |
| *Fos* | 0.914494 | -0.901906786 | 1 |
| *Egr1* | 0.020563 | -0.858840253 | 1 |
| *Cxcl1* | 0.095591 | -0.837843391 | 1 |
| *Tnc* | 0.005688 | -0.785981855 | 1 |
| *Igfbp5* | 0.152472 | -0.741471561 | 1 |
| *Trib1* | 0.004855 | -0.731839602 | 1 |
| *Gadd45g* | 0.096854 | -0.726784992 | 1 |

### Cluster 5 – Tip cells

#### Ranked by p value

| Gene | p_val | avg_log2FC | p_val_adj |
|---|---|---|---|
| *Nckap1* | 0.00064 | 0.63026736 | 1 |
| *Btnl9* | 0.001587 | 0.741823723 | 1 |
| *Wdr36* | 0.001589 | 0.478788519 | 1 |
| *Fnip1* | 0.001967 | 0.550363037 | 1 |
| *Armcx4* | 0.001987 | 0.439326755 | 1 |
| *C1galt1* | 0.00236 | 0.407872862 | 1 |
| *Tro* | 0.002515 | 0.615841405 | 1 |
| *Sidt2* | 0.002865 | 0.363869815 | 1 |
| *Lama4* | 0.003327 | 0.648724764 | 1 |
| *Agps* | 0.004004 | 0.459190136 | 1 |
| *Rps2* | 4.74E-06 | -0.443972202 | 0.153091 |
| *Rpl27* | 0.000417 | -0.349904108 | 1 |
| *Rpl30* | 0.001058 | -0.242215307 | 1 |
| *Gipc1* | 0.001315 | -0.620148394 | 1 |
| *Ndufa4* | 0.001355 | -0.529450154 | 1 |
| *Cd47* | 0.001356 | -0.521404893 | 1 |
| *Sh3gl3* | 0.001436 | -0.658069152 | 1 |
| *Rpl11* | 0.001496 | -0.294767887 | 1 |
| *Ubac2* | 0.00154 | -0.49390739 | 1 |
| *Rpp21* | 0.001649 | -0.630830191 | 1 |

#### Ranked by Log2FC

| Gene | p_val | avg_log2FC | p_val_adj |
|---|---|---|---|
| *Igfbp5* | 0.01801 | 0.981049397 | 1 |
| *Kdm6b* | 0.011774 | 0.804367729 | 1 |
| *Btnl9* | 0.001587 | 0.741823723 | 1 |
| *Dab2* | 0.034824 | 0.73791772 | 1 |
| *Cox19* | 0.266407 | 0.699966939 | 1 |
| *Ift122* | 0.183222 | 0.692890979 | 1 |
| *Atad2b* | 0.016298 | 0.676233777 | 1 |
| *Fn1* | 0.013995 | 0.671981039 | 1 |
| *Adamts9* | 0.028446 | 0.671640355 | 1 |
| *Nsrp1* | 0.077285 | 0.667201968 | 1 |
| *Xist* | 0.09093 | -0.747910089 | 1 |
| *Sde2* | 0.020343 | -0.720800367 | 1 |
| *Sh3gl3* | 0.001436 | -0.658069152 | 1 |
| *Zfp512* | 0.644441 | -0.643466585 | 1 |
| *Rpp21* | 0.001649 | -0.630830191 | 1 |
| *Gipc1* | 0.001315 | -0.620148394 | 1 |
| *Ece1* | 0.003257 | -0.611939817 | 1 |
| *Ppp1r11* | 0.002116 | -0.606942239 | 1 |
| *Fabp4* | 0.045185 | -0.599863737 | 1 |
| *Odc1* | 0.083285 | -0.598406728 | 1 |

### Cluster 6 – Venous ECs

#### Ranked by p value

| Gene | p_val | avg_log2FC | p_val_adj |
|---|---|---|---|
| ***Wtip*** | 0.00029 | 0.790020353 | 1 |
| ***Agrn*** | 0.000664 | 1.064052124 | 1 |
| ***Nsd3*** | 0.001013 | 0.874296003 | 1 |
| ***Pak1ip1*** | 0.001194 | 0.751742648 | 1 |
| ***Diaph1*** | 0.001209 | 0.640771408 | 1 |
| ***Trappc6b*** | 0.001357 | 0.672180863 | 1 |
| ***Gna12*** | 0.001411 | 0.620251361 | 1 |
| ***Ptprg*** | 0.001904 | 0.720076814 | 1 |
| ***Parvb*** | 0.002228 | 0.572123339 | 1 |
| ***Ctdnep1*** | 0.002331 | 0.669643628 | 1 |
| ***Rps12*** | 0.002144 | -0.334220978 | 1 |
| ***Rps2*** | 0.002436 | -0.658655664 | 1 |
| ***Gpr180*** | 0.0085 | -0.754015522 | 1 |
| ***Tjp1*** | 0.009014 | -0.780757121 | 1 |
| ***Psmb3*** | 0.010079 | -0.774203726 | 1 |
| ***Snrpe*** | 0.010141 | -0.593443475 | 1 |
| ***Hnrnpa2b1*** | 0.011329 | -0.468482407 | 1 |
| ***Hmgb2*** | 0.012615 | -0.838513448 | 1 |
| ***Rplp2*** | 0.012631 | -0.337848501 | 1 |
| ***Nudt4*** | 0.014513 | -0.720094626 | 1 |

#### Ranked by Log2FC

| Gene | p_val | avg_log2FC | p_val_adj |
|---|---|---|---|
| ***Ccn2*** | 0.058643 | 1.428921609 | 1 |
| ***Peg3*** | 0.112643 | 1.33874735 | 1 |
| ***Igf2r*** | 0.015921 | 1.115063762 | 1 |
| ***Sox17*** | 0.771611 | 1.083109485 | 1 |
| ***Igf1*** | 0.010474 | 1.072954376 | 1 |
| ***Cdkn1c*** | 0.141857 | 1.066965392 | 1 |
| ***Agrn*** | 0.000664 | 1.064052124 | 1 |
| ***Cttnbp2nl*** | 0.006032 | 1.044407039 | 1 |
| ***Igfbp5*** | 0.274563 | 1.041370175 | 1 |
| ***Socs2*** | 0.042166 | 0.977220331 | 1 |
| ***Fos*** | 0.762667 | -1.892962345 | 1 |
| ***Xist*** | 0.071594 | -0.910318708 | 1 |
| ***Mrpl18*** | 0.133962 | -0.892184712 | 1 |
| ***Hmgb2*** | 0.012615 | -0.838513448 | 1 |
| ***Itch*** | 0.030453 | -0.824762922 | 1 |
| ***Smc3*** | 0.049188 | -0.814839112 | 1 |
| ***Tjp1*** | 0.009014 | -0.780757121 | 1 |
| ***Npr3*** | 0.166771 | -0.779990691 | 1 |
| ***Ggta1*** | 0.282903 | -0.77522344 | 1 |
| ***Psmb3*** | 0.010079 | -0.774203726 | 1 |

**Supplemental Figure S1: Nkx2.5-CRE activity in the early embryo**

Representative *Nkx2.5-CRE;ROSA26-lacZ* embryos indicate the activity of *Nkx2.5-CRE* (detected by blue X-Gal stain) in the heart during embryonic development Scale bar = 1mm. **A-B** wholemount embryos, **C** wholemount heart, **D** sagittal section and **E** transverse section. Age indicated in image, ht = heart; ra = right atrium; la = left atrium; rv = right ventricle; lv = left ventricle; oft = outflow tract.

Nkx2.5-CRE; Rosa*LacZ*

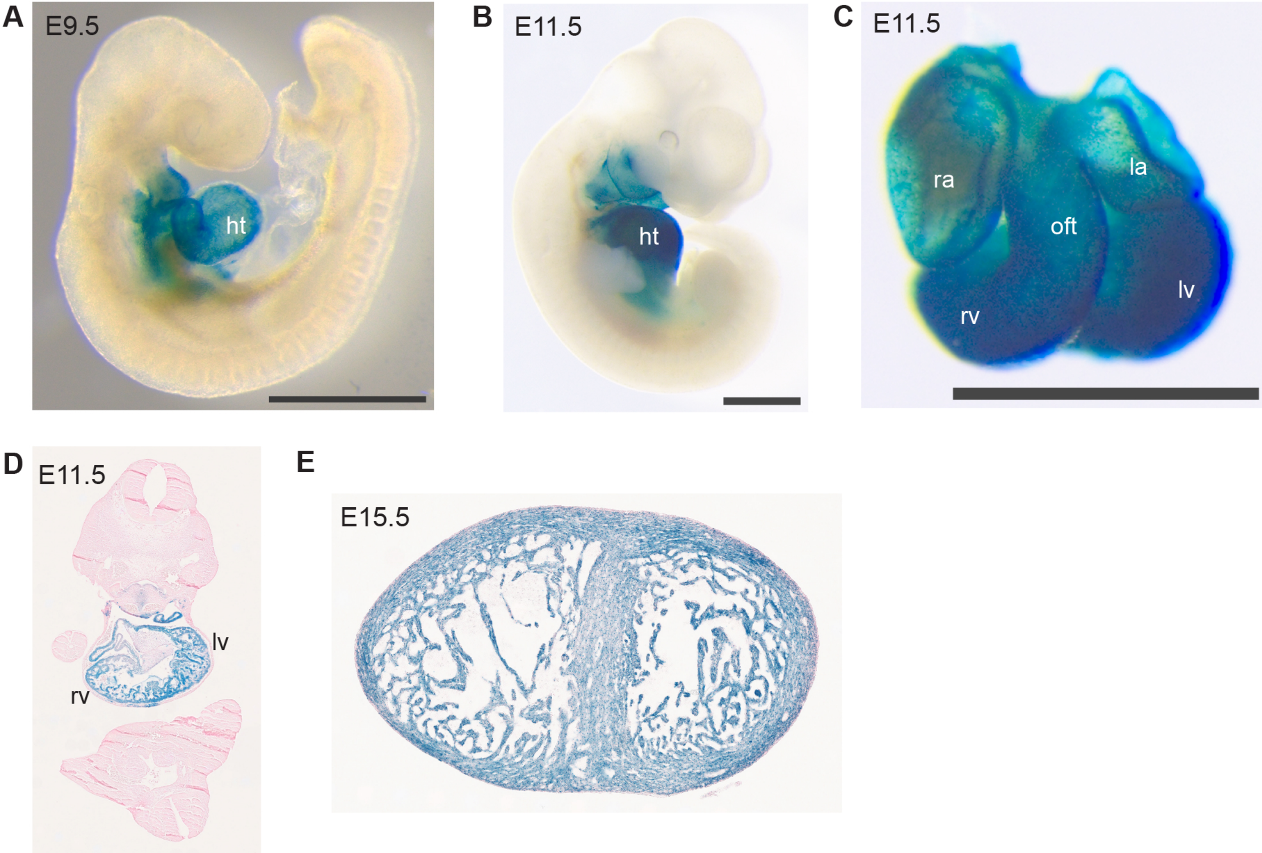

**Supplemental Figure S2: Vascular development in hypoxic myocardium**

Dorsal (**A-B**) and ventral (**C**) views of E12.5 (**A**) and E13.5 (**B-C**) *Phd2*$^{fl/fl}$ and *Phd2*$^{fl/fl}$*;Nkx2.5-Cre* hearts immunostained for the endothelial marker CD31 , see also Figure 1H. **D** Quantification of the extension of the SV-derived vascular plexus as a percentage of the total dorsal area of E13.5 hearts. p=0.0535 (unpaired *t* test), error bars show SEM, N=2 hearts per genotype. **E** Quantification of the number of blood islands on the ventral aspect of E13.5 *Phd2*$^{fl/fl}$ and *Phd2*$^{fl/fl}$*;Nkx2.5-Cre* hearts. NS = not significant (unpaired *t* test), error bars show SEM, N=5 *Phd2*$^{fl/fl}$*;Nkx2.5-Cre* hearts and N= 7 *Phd2*$^{fl/fl}$ hearts.

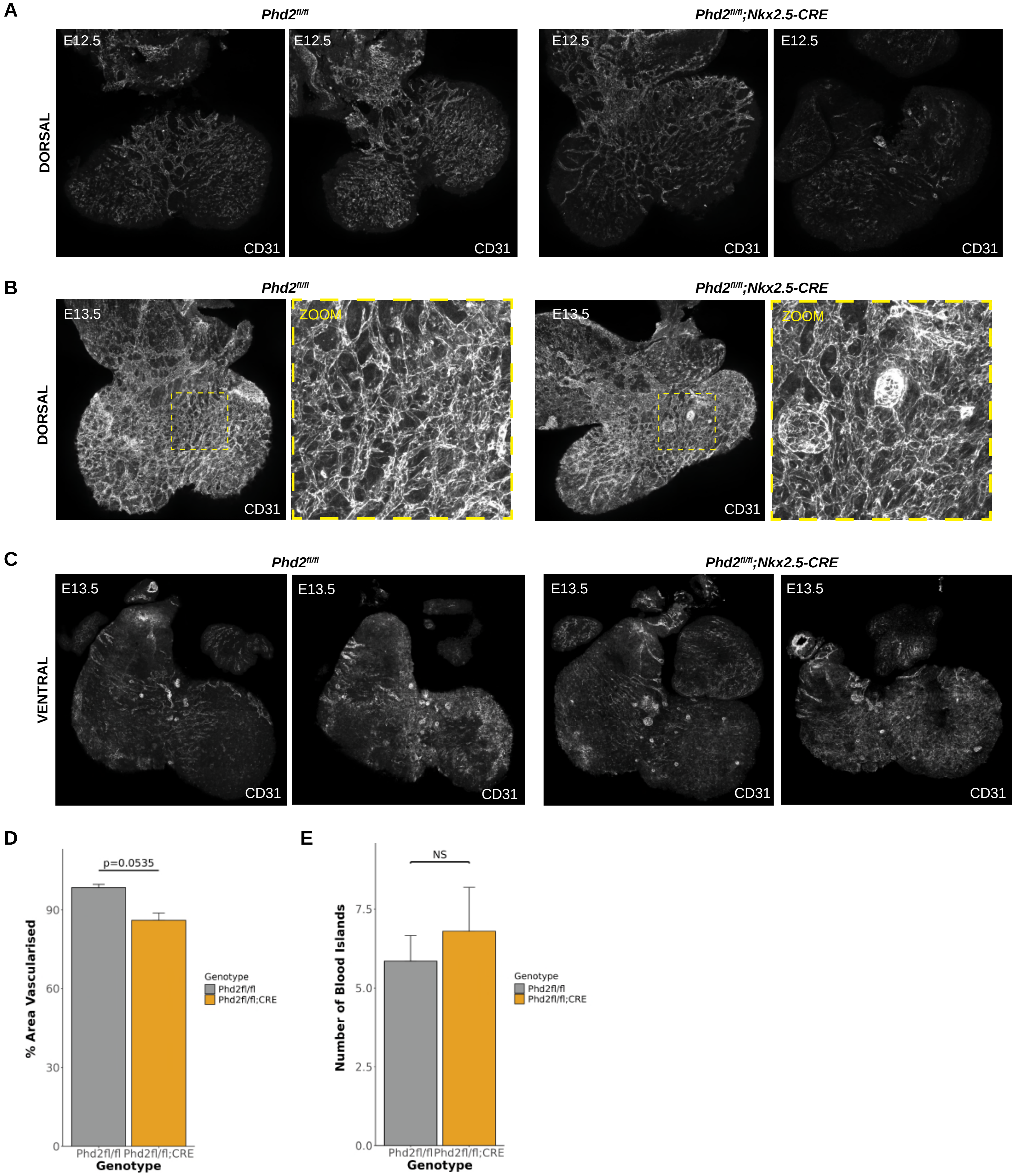

A
Phd2fl/fl
Phd2fl/fl;Nkx2.5-CRE
DORSAL
E12.5
CD31
B
E13.5
ZOOM
C
VENTRAL
D
p=0.0535
% Area Vascularised
Genotype
Phd2fl/fl
Phd2fl/fl;CRE
E
NS
Number of Blood Islands

**Supplemental Figure S3: Quality control for sc-RNAseq dataset**

**A** Sequencing statistics for the *Phd2*$^{fl/fl}$ and *Phd2*$^{fl/fl}$*;Nkx2.5-CRE* groups of cells. **B** Number of reads per cell, number of genes expressed per cell, and the percentage of genes mapped to the mitochondrial genome, shown per cluster for all the cells collected. **C** Number of reads per cell, number of genes expressed per cell, and the percentage of mitochondrial genes shown per cluster for the coronary EC-only dataset.

A

| | *Phd2$^{fl/fl}$* | *Phd2$^{fl/fl}$;Nkx2.5-Cre* |
|---|---|---|
| **Sequencing** | | |
| Number of Reads | 349950236 | 296927485 |
| Valid Barcodes | 96.9% | 96.7% |
| Sequencing Saturation | 53.3% | 50.3% |
| Q30 Bases in Barcode | 96.9% | 96.9% |
| Q30 Bases in RNA Read | 95.7% | 95.7% |
| Q30 Bases in UMI | 97.2% | 97.2% |
| **Mapping** | | |
| Reads Mapped to Genome | 94.7% | 94.2% |
| Reads Mapped Confidently to Genome | 91.6% | 91.1% |
| Reads Mapped Confidently to Intergenic Regions | 6.1% | 6.0% |
| Reads Mapped Confidently to Intronic Regions | 31.3% | 31.4% |
| Reads Mapped Confidently to Exonic Regions | 54.2% | 53.6% |
| Reads Mapped Confidently to Transcriptome | 49.5% | 49.0% |
| Reads Mapped Antisense to Gene | 3.6% | 3.5% |
| **Cells** | | |
| Estimated Number of Cells | 5403 | 3200 |
| Fraction Reads in Cells | 91.7% | 90.7% |
| Mean Reads per Cell | 64770 | 92790 |
| Median Genes per Cell | 3538 | 4620 |
| Total Genes Detected | 22680 | 22282 |
| Median UMI Counts per Cell | 10842 | 17678 |

B

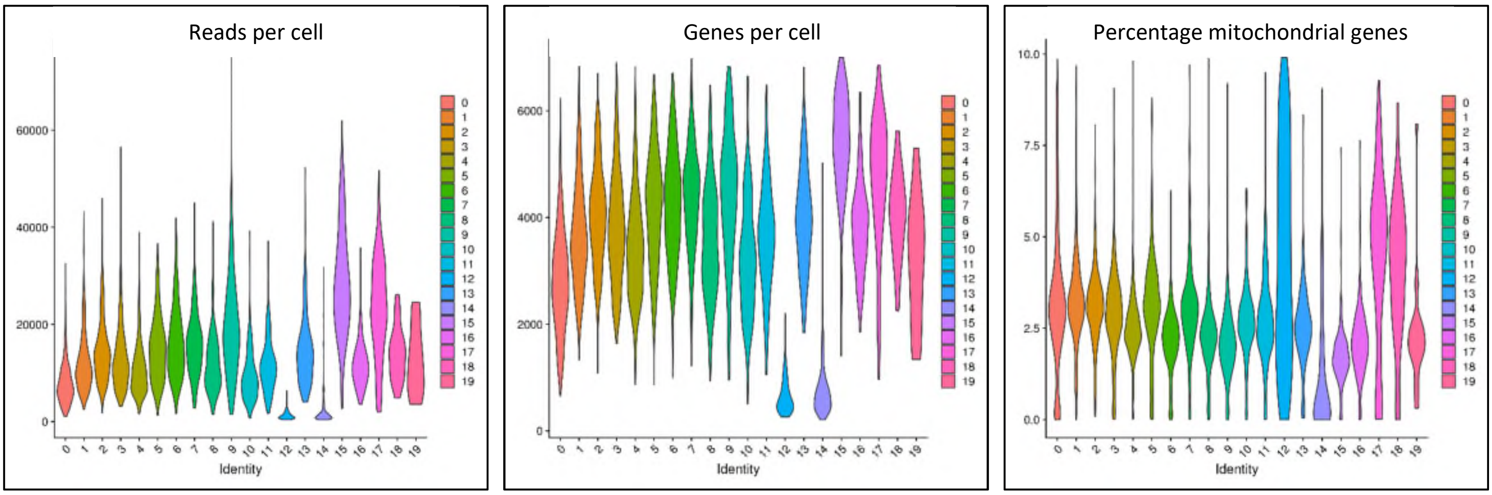

C

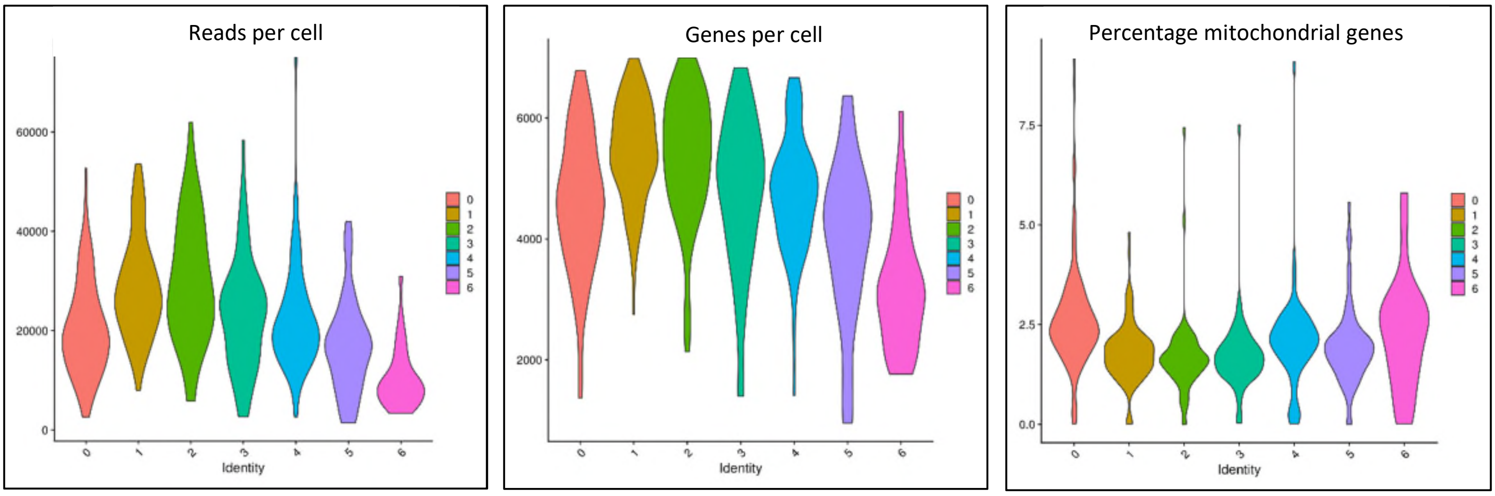

**Supplemental Figure S4: SOX17 expression in the developing coronary vasculature**

**A** Ventral and dorsal views of the E12.5 heart with immunostaining for SOX17 protein and the endothelial marker Endomucin (EMCN). Magnified views show nuclear SOX17 throughout the developing vasculature and within round blood islands on the ventral aspect. **B** SOX17 and EMCN immunostaining of the E13.5 heart.

All scale bars = 0.2mm.

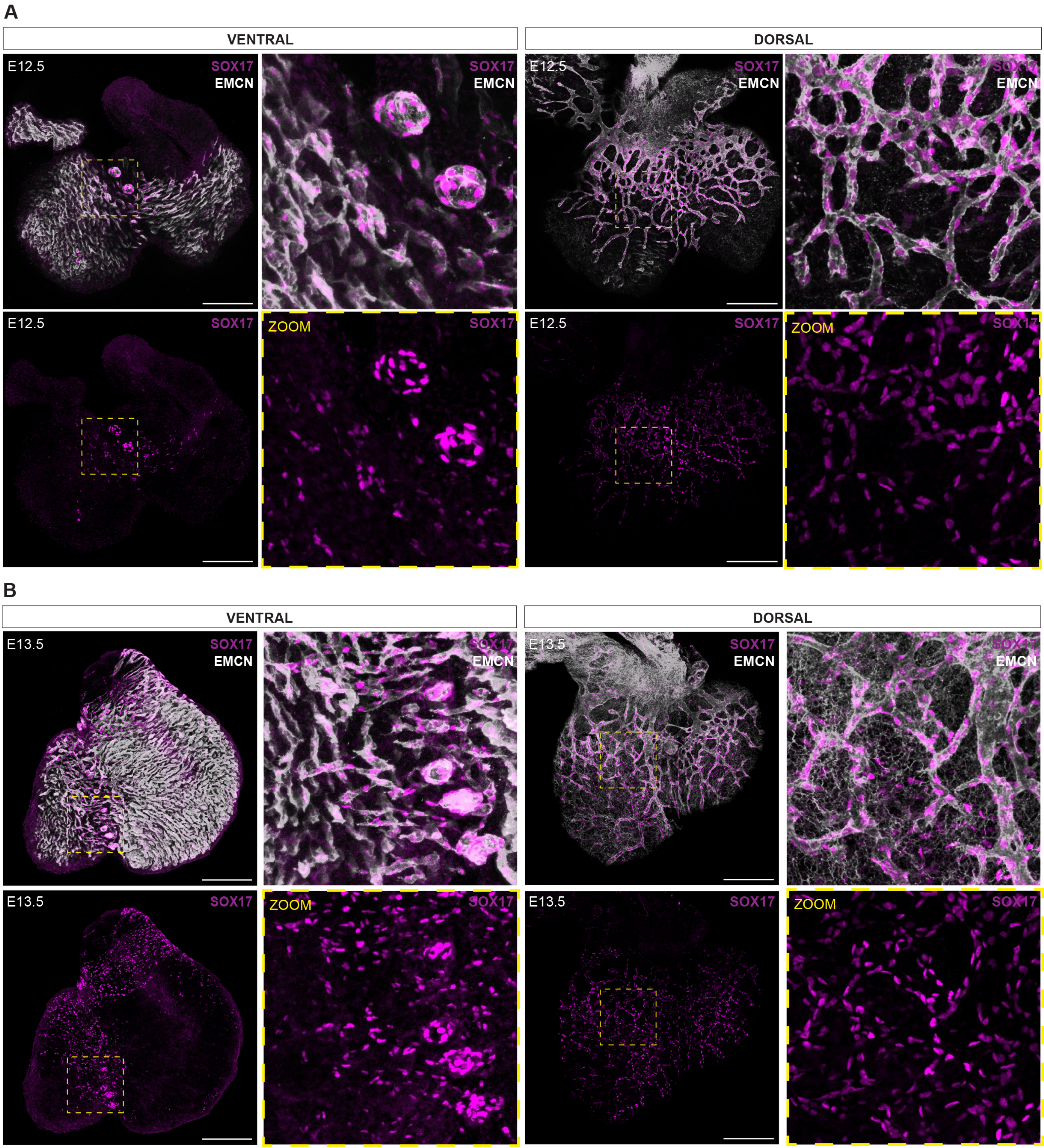
A
VENTRAL
DORSAL
E12.5
SOX17
EMCN
ZOOM
B
E13.5

**Supplemental Figure S5: E11.75 and E12.5 HLX-3:*LacZ* hearts**

**A** Additional *Phd2*$^{fl/fl}$ and *Phd2*$^{fl/fl}$*;Nkx2.5-CRE* hearts collected at E11.75 expressing the HLX-3:*LacZ* transgene, collected across two litters. The hearts shown in Figure 4C were also from Litter 1. **B** Two additional litters of *Phd2*$^{fl/fl}$ and *Phd2*$^{fl/fl}$*;Nkx2.5-CRE* E12.5 hearts. The hearts in Litter 2 were stained for longer than other hearts, so the blue XGal stain is stronger.

All scale bars = 1mm.

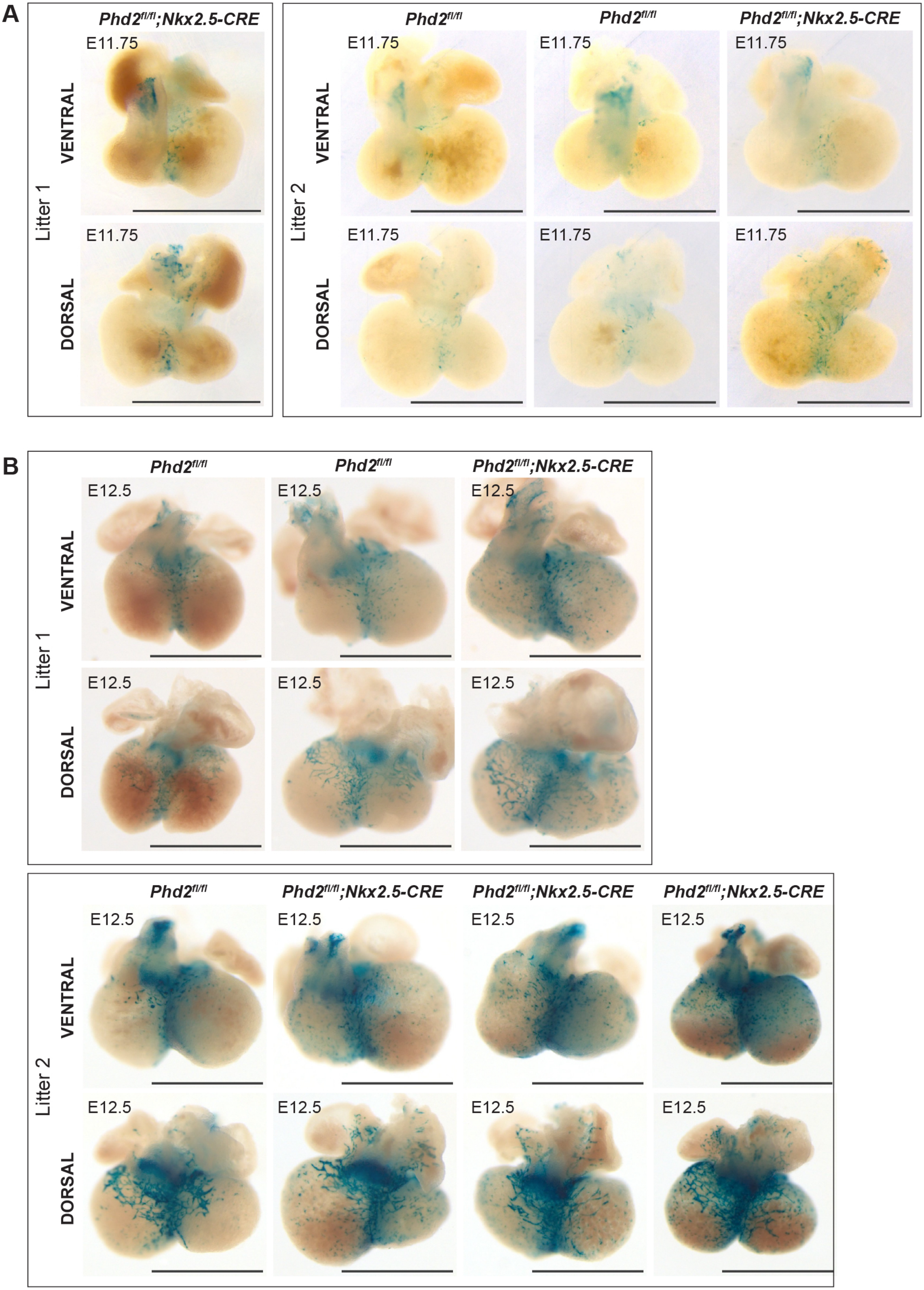
A
Phd2fl/fl;Nkx2.5-CRE
Phd2fl/fl
Phd2fl/fl
Phd2fl/fl;Nkx2.5-CRE
E11.75
VENTRAL
DORSAL
Litter 1
Litter 2
B
Phd2fl/fl
Phd2fl/fl
Phd2fl/fl;Nkx2.5-CRE
E12.5
VENTRAL
DORSAL
Litter 1
Phd2fl/fl
Phd2fl/fl;Nkx2.5-CRE
Phd2fl/fl;Nkx2.5-CRE
Phd2fl/fl;Nkx2.5-CRE
Litter 2

**Supplemental Figure S6: E13.5 HLX-3:*LacZ* hearts**

3 litters of *Phd2*$^{fl/fl}$ and *Phd2*$^{fl/fl}$*;Nkx2.5-CRE* hearts collected at E13.5 expressing the HLX-3:*LacZ* transgene, in addition to the littermate hearts shows in Figure 4F. Scale bars = 1mm.

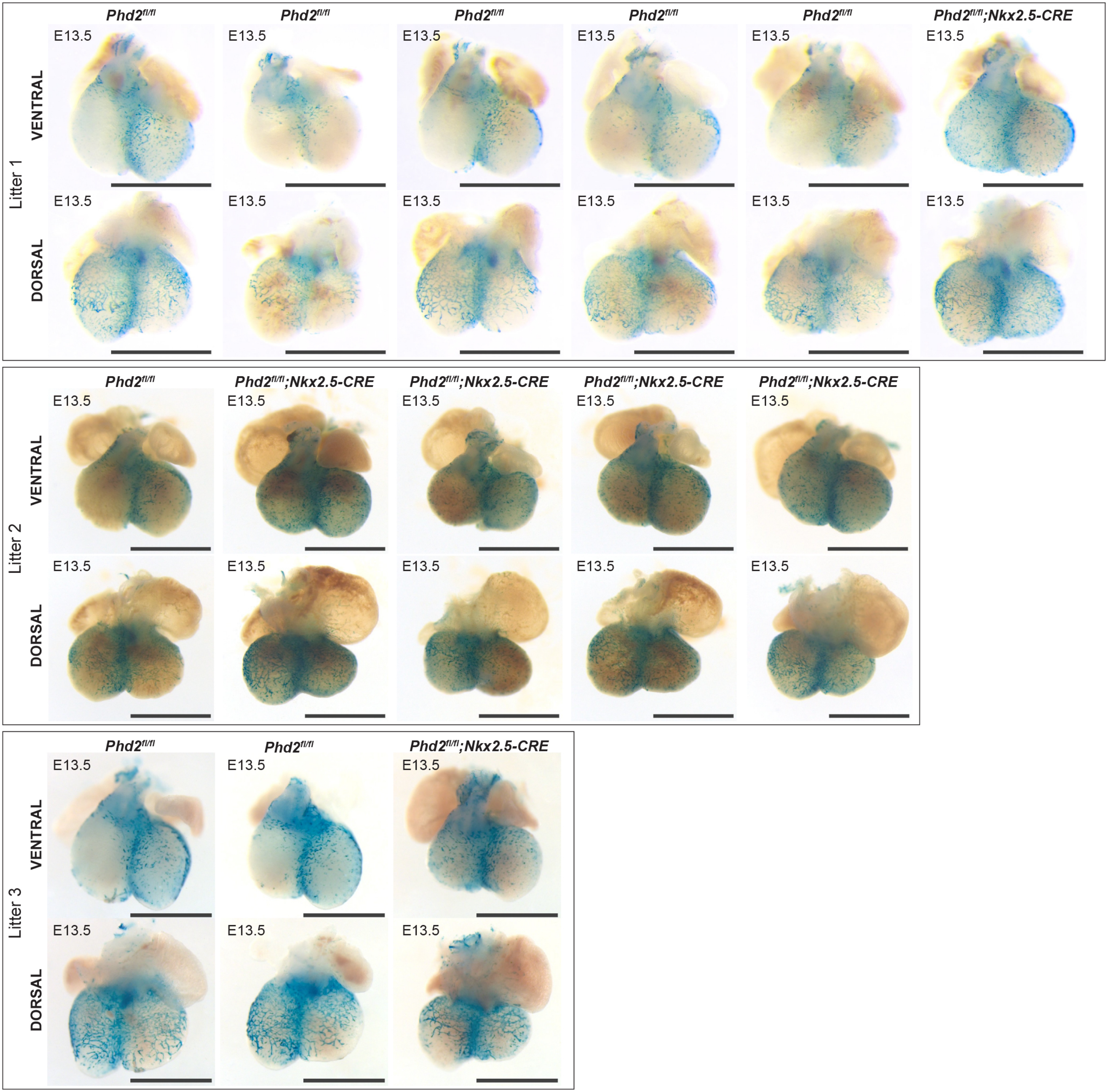

Litter 1
VENTRAL
DORSAL
Phd2fl/fl
Phd2fl/fl
Phd2fl/fl
Phd2fl/fl
Phd2fl/fl
Phd2fl/fl;Nkx2.5-CRE
E13.5
Litter 2
Phd2fl/fl
Phd2fl/fl;Nkx2.5-CRE
Phd2fl/fl;Nkx2.5-CRE
Phd2fl/fl;Nkx2.5-CRE
Phd2fl/fl;Nkx2.5-CRE
Litter 3
Phd2fl/fl
Phd2fl/fl
Phd2fl/fl;Nkx2.5-CRE

**Supplemental Figure S7: E15.5 HLX-3:*LacZ* hearts**

**A** All additional hearts collected at E15.5 across 5 litters, hearts shown in main Figure 4I were from Litter 1. Litter 5 contained 3 heterozygous *Phd2*$^{fl/+}$*;Nkx2.5-CRE* hearts, whose expression is consistent with the *Phd2*$^{fl/fl}$ hearts and therefore show that Cre activity is not affecting HLX-3:*LacZ* activity. Scale bars = 1mm. **B** Transverse sections through a *Phd2*$^{fl/fl}$ and *Phd2*$^{fl/fl}$*;Nkx2.5-CRE* heart, with magnified image of the interventricular septum (IVS). Scale bars = 0.2mm.

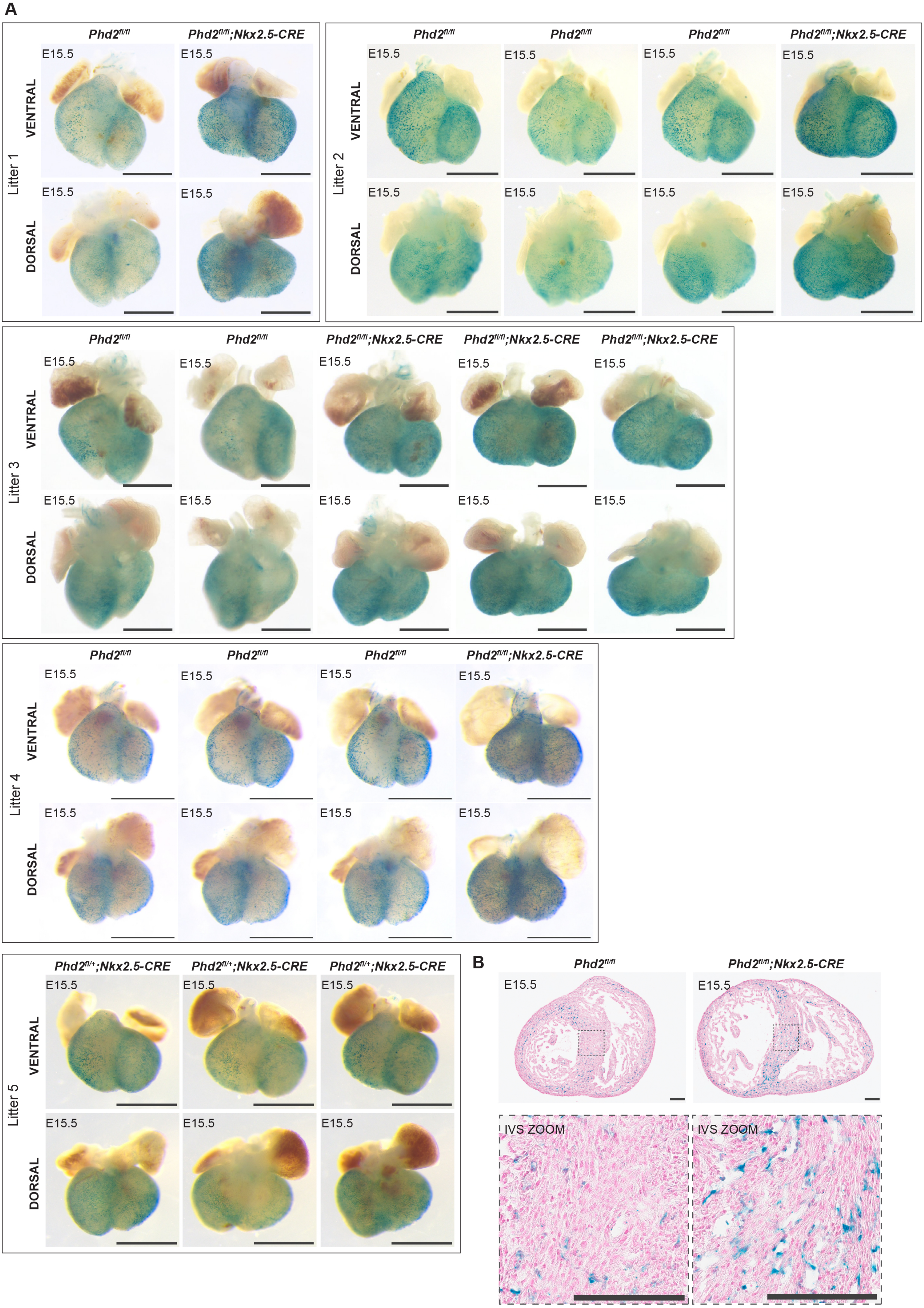

A
Litter 1
Phd2fl/fl
Phd2fl/fl;Nkx2.5-CRE
VENTRAL
DORSAL
E15.5
Litter 2
Phd2fl/fl
Phd2fl/fl
Phd2fl/fl
Phd2fl/fl;Nkx2.5-CRE
Litter 3
Phd2fl/fl
Phd2fl/fl
Phd2fl/fl;Nkx2.5-CRE
Phd2fl/fl;Nkx2.5-CRE
Phd2fl/fl;Nkx2.5-CRE
Litter 4
Phd2fl/fl
Phd2fl/fl
Phd2fl/fl
Phd2fl/fl;Nkx2.5-CRE
Litter 5
Phd2fl/+;Nkx2.5-CRE
Phd2fl/+;Nkx2.5-CRE
Phd2fl/+;Nkx2.5-CRE
B
Phd2fl/fl
Phd2fl/fl;Nkx2.5-CRE
IVS ZOOM

**Supplemental Figure S8: E12.5 and E13.5 NOTCH1+16:*LacZ* hearts**

**A** Additional *Phd2*$^{fl/fl}$ and *Phd2*$^{fl/fl}$*;Nkx2.5-CRE* hearts collected at E12.5 expressing the NOTCH1+16:*LacZ* transgene, collected across four litters. Litter 1 also contained the two hearts shown in Figure 5B. Scale bars = 1mm. **B** Additional hearts collected at E13.5 expressing the NOTCH1+16:*LacZ* transgene. The two hearts shown in Figure 5C were from Litter 1. Scale bars = 1mm.

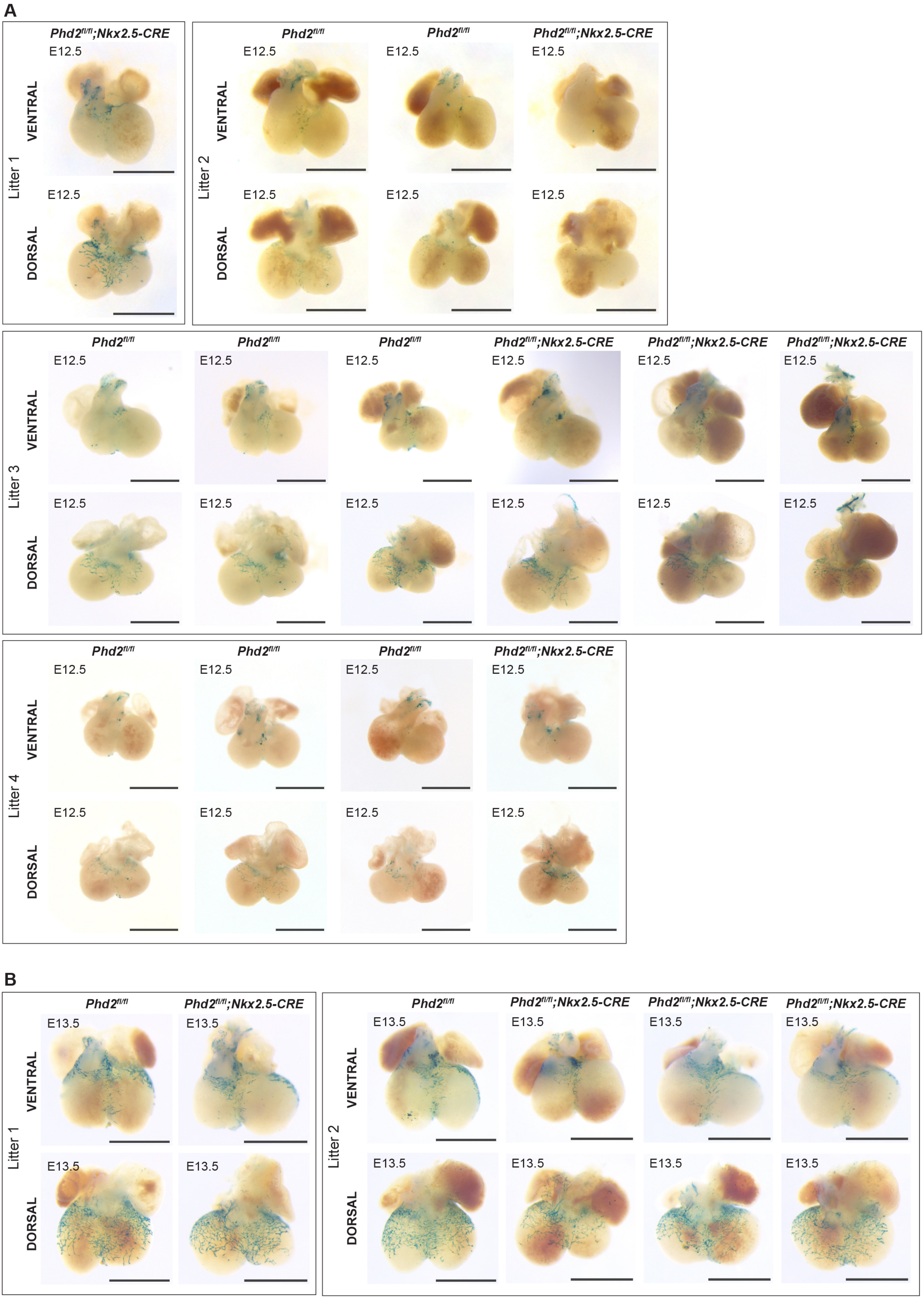

A
Litter 1
Litter 2
Litter 3
Litter 4
VENTRAL
DORSAL
$Phd2^{fl/fl}$
$Phd2^{fl/fl}$;Nkx2.5-CRE
E12.5
B
E13.5

**Supplemental Figure S9: E15.5 NOTCH1+16:*LacZ* hearts**

Additional *Phd2*$^{fl/fl}$ and *Phd2*$^{fl/fl}$*;Nkx2.5-CRE* hearts expressing the NOTCH1+16:*LacZ* transgene, collected at E15.5. Litter 1 also contained the hearts shown in Figure 5F. Scale bars = 1mm.

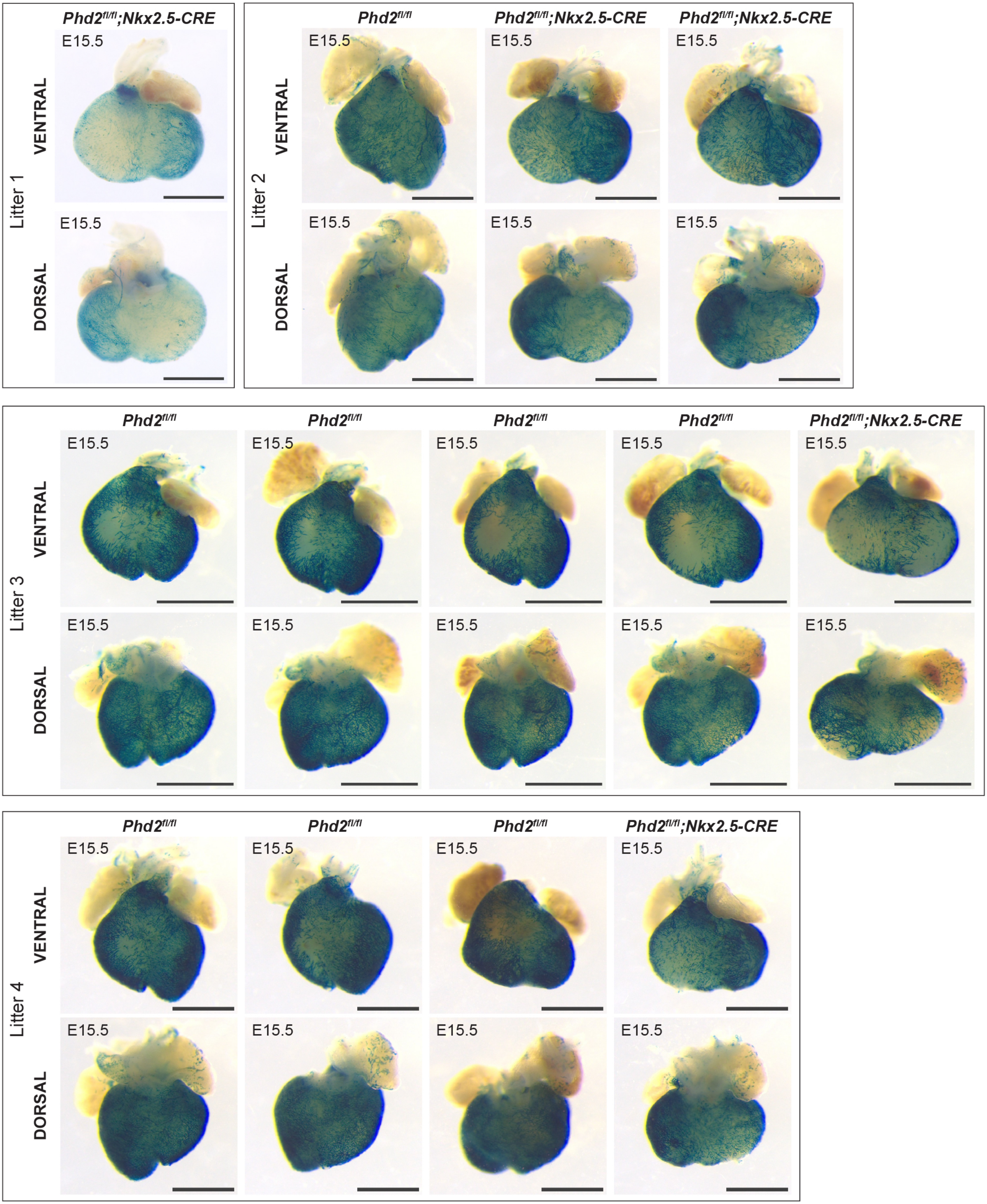

Litter 1
Phd2fl/fl;Nkx2.5-CRE
VENTRAL
DORSAL
E15.5
Litter 2
Phd2fl/fl
Phd2fl/fl;Nkx2.5-CRE
Phd2fl/fl;Nkx2.5-CRE
Litter 3
Phd2fl/fl
Phd2fl/fl
Phd2fl/fl
Phd2fl/fl
Phd2fl/fl;Nkx2.5-CRE
Litter 4
Phd2fl/fl
Phd2fl/fl
Phd2fl/fl
Phd2fl/fl;Nkx2.5-CRE

**Supplemental Figure S10: E13.5 Dll4-12:*LacZ* hearts**

Additional *Phd2*$^{fl/fl}$ and *Phd2*$^{fl/fl}$*;Nkx2.5-CRE* hearts expressing the Dll4-12:*LacZ* transgene, collected at E13.5. The hearts shown in main Figure 6B were also from Litter 1. Scale bars = 1mm.

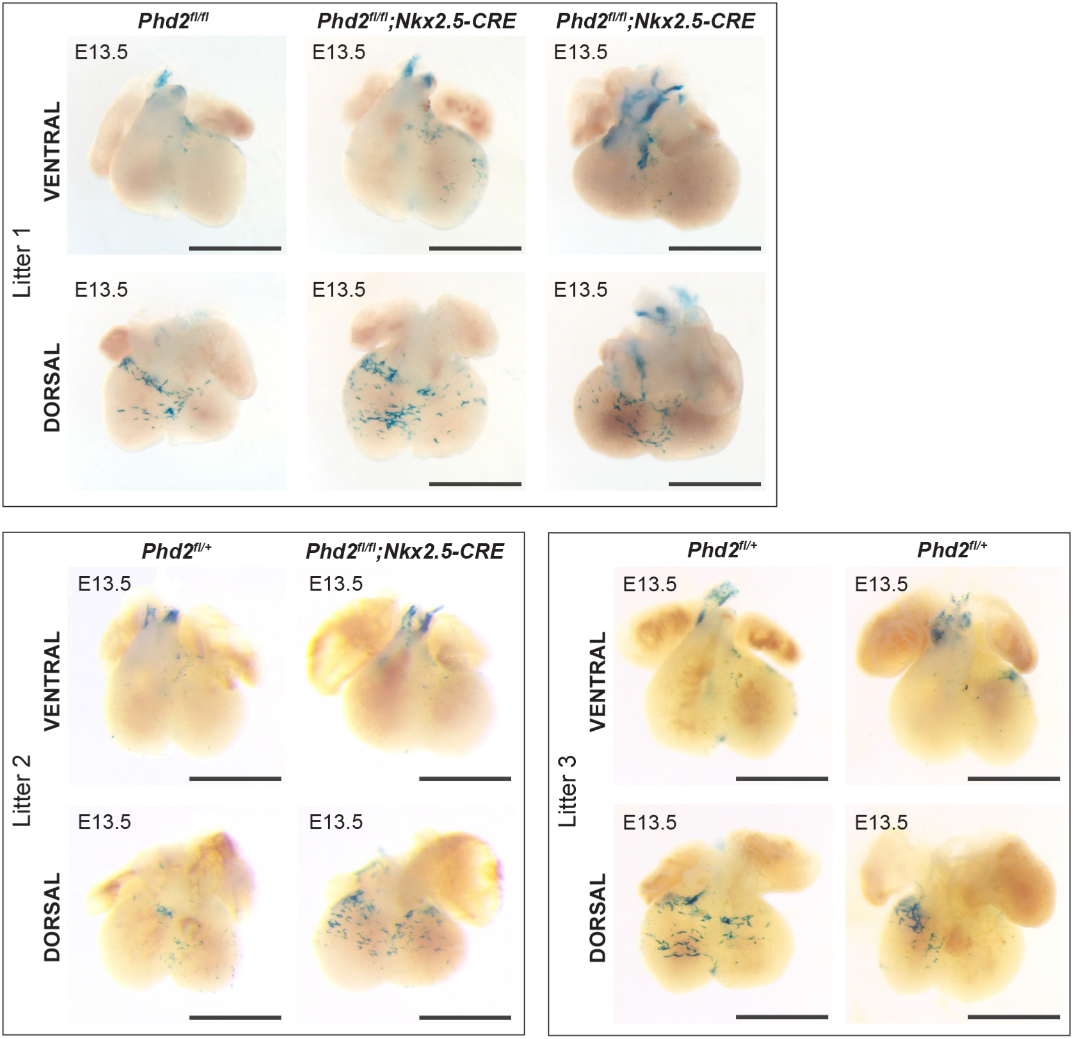
Litter 1
$Phd2^{fl/fl}$
$Phd2^{fl/fl}$;Nkx2.5-CRE
$Phd2^{fl/fl}$;Nkx2.5-CRE
VENTRAL
DORSAL
E13.5
Litter 2
$Phd2^{fl/+}$
$Phd2^{fl/fl}$;Nkx2.5-CRE
Litter 3
$Phd2^{fl/+}$
$Phd2^{fl/+}$

**Supplemental Figure S11: E15.5 Dll4-12:*LacZ* hearts**

Additional 5 litters of E15.5 hearts to those shown in Figure 6D, showing expression of the Dll4-12:*LacZ* transgene on different genetic backgrounds. The heterozygous *Phd2*$^{fl/+}$*;Nkx2.5-CRE* hearts show more similar activity to their CRE-negative littermate hearts, and therefore indicate there is no effect of CRE expression on the enhancer activity. Black arrows indicate left anterior descending artery. Scale bars = 1mm.

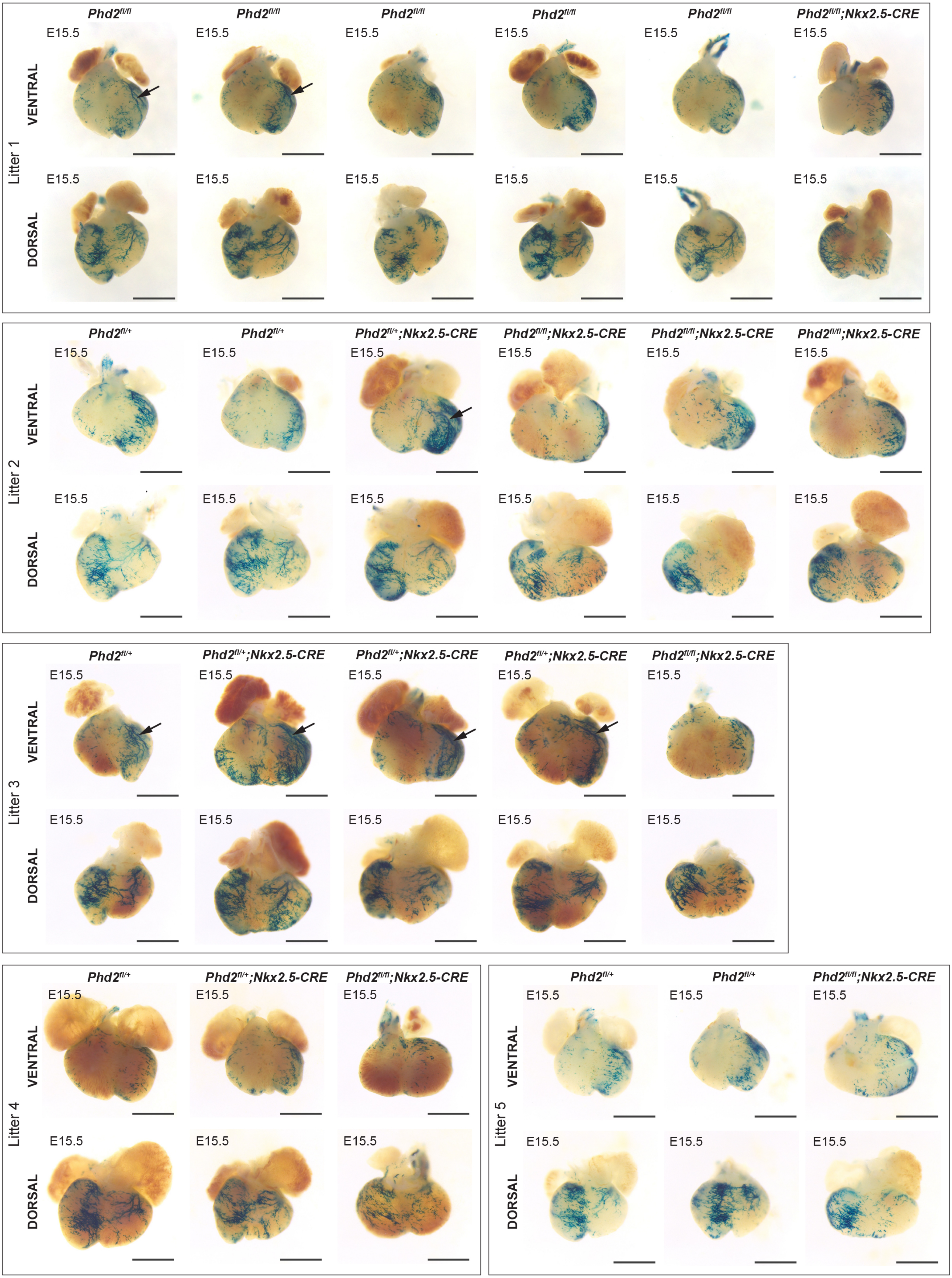

Litter 1
VENTRAL
DORSAL
$Phd2^{fl/fl}$
$Phd2^{fl/fl}$
$Phd2^{fl/fl}$
$Phd2^{fl/fl}$
$Phd2^{fl/fl}$
$Phd2^{fl/fl}$;Nkx2.5-CRE
E15.5
Litter 2
VENTRAL
DORSAL
$Phd2^{fl/+}$
$Phd2^{fl/+}$
$Phd2^{fl/+}$;Nkx2.5-CRE
$Phd2^{fl/fl}$;Nkx2.5-CRE
$Phd2^{fl/fl}$;Nkx2.5-CRE
$Phd2^{fl/fl}$;Nkx2.5-CRE
E15.5
Litter 3
VENTRAL
DORSAL
$Phd2^{fl/+}$
$Phd2^{fl/+}$;Nkx2.5-CRE
$Phd2^{fl/+}$;Nkx2.5-CRE
$Phd2^{fl/+}$;Nkx2.5-CRE
$Phd2^{fl/fl}$;Nkx2.5-CRE
E15.5
Litter 4
VENTRAL
DORSAL
$Phd2^{fl/+}$
$Phd2^{fl/+}$;Nkx2.5-CRE
$Phd2^{fl/fl}$;Nkx2.5-CRE
E15.5
Litter 5
VENTRAL
DORSAL
$Phd2^{fl/+}$
$Phd2^{fl/+}$
$Phd2^{fl/fl}$;Nkx2.5-CRE
E15.5

**Supplemental Figure S12: E18.5 and P6-8 Dll4-12:*LacZ* hearts**

**A** 3 additional litters of E18.5 hearts to those shown in Figure 6D, showing expression of the Dll4-12:*LacZ* transgene on different genetic backgrounds. The heterozygous *Phd2*$^{fl/+}$*;Nkx2.5-CRE* hearts are controls for CRE activity. Black arrows indicate left anterior descending artery. Scale bars = 1mm. **B** Postnatal activity of Dll4-12:*LacZ* in P6-P8 hearts, in addition to those in Figure 6F, shows similar activity in hypoxic myocardium and controls. Black arrows indicate left anterior descending artery. Scale bars = 1mm.

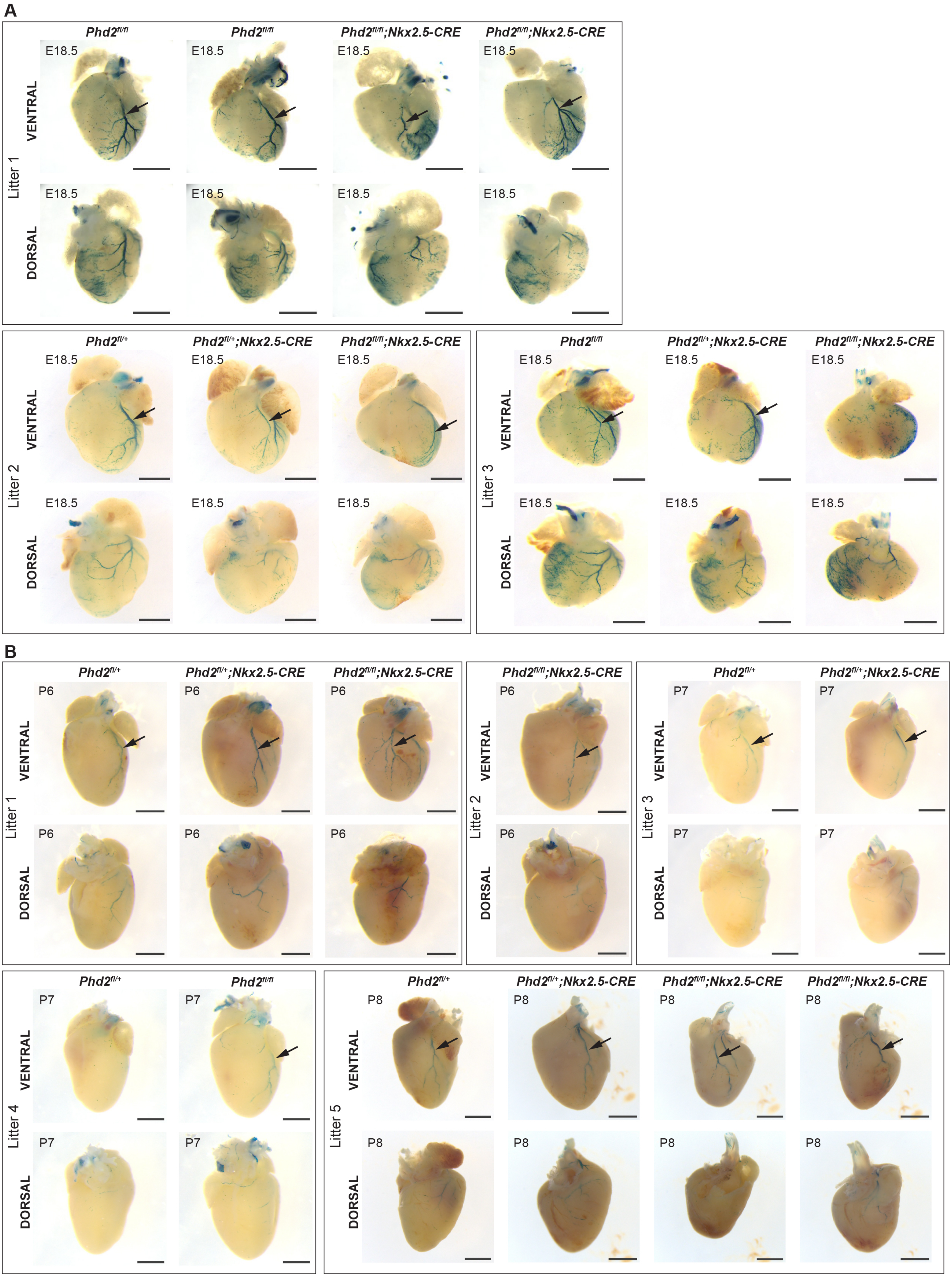
A
Phd2fl/fl
Phd2fl/fl
Phd2fl/fl;Nkx2.5-CRE
Phd2fl/fl;Nkx2.5-CRE
E18.5
VENTRAL
DORSAL
Litter 1
Phd2fl/+
Phd2fl/+;Nkx2.5-CRE
Phd2fl/fl;Nkx2.5-CRE
Litter 2
Phd2fl/fl
Phd2fl/+;Nkx2.5-CRE
Phd2fl/fl;Nkx2.5-CRE
Litter 3
B
Phd2fl/+
Phd2fl/+;Nkx2.5-CRE
Phd2fl/fl;Nkx2.5-CRE
P6
Litter 1
Phd2fl/fl;Nkx2.5-CRE
Litter 2
Phd2fl/+
Phd2fl/+;Nkx2.5-CRE
P7
Litter 3
Phd2fl/+
Phd2fl/fl
Litter 4
Phd2fl/+
Phd2fl/+;Nkx2.5-CRE
Phd2fl/fl;Nkx2.5-CRE
Phd2fl/fl;Nkx2.5-CRE
P8
Litter 5

**Supplemental Figure S13: E13.5 and E15.5 EphB4-2:*LacZ* hearts**

**A** E13.5 hearts showing expression of the EphB4-2:*LacZ* transgene on *Phd2*$^{fl/fl}$ and *Phd2*$^{fl/fl}$*;Nkx2.5-CRE* backgrounds, in addition to those shown in Figure 7B,. The hearts in Figure 7B were also from Litter 1. Scale bars = 1mm. **B** Activity of EphB4-2:*LacZ* in E15.5 hearts, in addition to those in Figure 7D. Scale bars = 1mm.

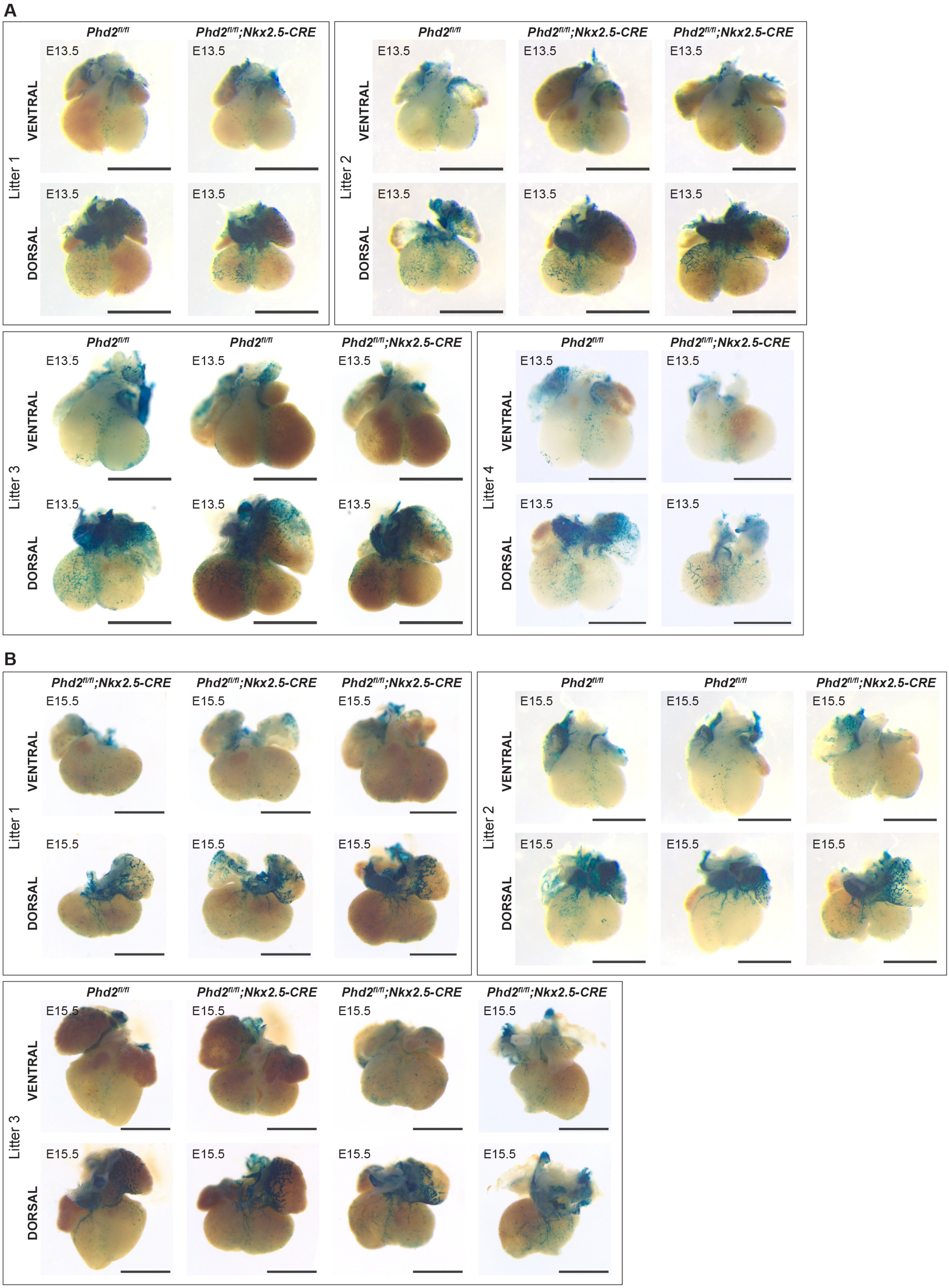
A
Litter 1
Phd2fl/fl
Phd2fl/fl;Nkx2.5-CRE
VENTRAL
DORSAL
E13.5
Litter 2
Phd2fl/fl
Phd2fl/fl;Nkx2.5-CRE
Phd2fl/fl;Nkx2.5-CRE
Litter 3
Phd2fl/fl
Phd2fl/fl
Phd2fl/fl;Nkx2.5-CRE
Litter 4
Phd2fl/fl
Phd2fl/fl;Nkx2.5-CRE
B
Litter 1
Phd2fl/fl;Nkx2.5-CRE
Phd2fl/fl;Nkx2.5-CRE
Phd2fl/fl;Nkx2.5-CRE
E15.5
Litter 2
Phd2fl/fl
Phd2fl/fl
Phd2fl/fl;Nkx2.5-CRE
Litter 3
Phd2fl/fl
Phd2fl/fl;Nkx2.5-CRE
Phd2fl/fl;Nkx2.5-CRE
Phd2fl/fl;Nkx2.5-CRE

**Supplemental Figure S14: E18.5 EphB4-2:*LacZ* hearts**

Additional *Phd2*$^{fl/fl}$ and *Phd2*$^{fl/fl}$*;Nkx2.5-CRE* hearts expressing the EphB4-2:*LacZ* transgene, collected at E18.5. Litter 1 also contained the hearts shown in Figure 7E. Scale bars = 1mm.

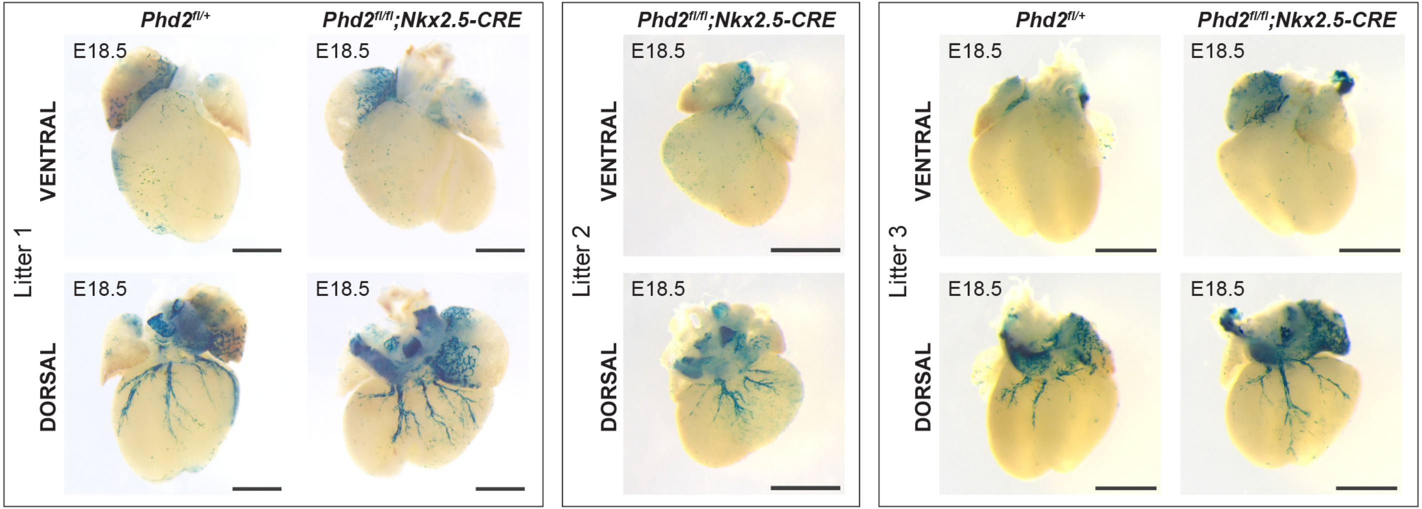

Phd2fl/+
Phd2fl/fl;Nkx2.5-CRE
Phd2fl/fl;Nkx2.5-CRE
Phd2fl/+
Phd2fl/fl;Nkx2.5-CRE
E18.5
VENTRAL
DORSAL
Litter 1
Litter 2
Litter 3
